# 3-D Ontogenetic Staging Atlas of the Epaulette Shark *Hemiscyllium ocellatum,* a Laboratory Model for Shark Development

**DOI:** 10.64898/2026.04.27.721166

**Authors:** Rebecca Dale, Frank Tulenko, Lucy Hersey, Peter Currie

**Author notes:** Authors contributed equally (Rebecca Dale, Frank Tulenko).

## Abstract

Chondrichthyans (cartilaginous fishes) form the sister group to osteichthyans (bony fishes) and therefore occupy a key phylogenetic position for comparative studies of early vertebrate evolution. Despite their importance, chondrichthyan development remains understudied relative to established model systems such as mouse, chick, and zebrafish, in part because of limited embryo accessibility and the lack of standardized laboratory resources for rearing. Here, we present the epaulette shark *Hemiscyllium ocellatum*, a small, oviparous shark as a tractable laboratory system for studying shark development. We provide an overview of epaulette shark husbandry requirements and generate a comprehensive micro-computed tomography imaging series spanning embryonic development through hatching. This dataset provides a three-dimensional anatomical atlas of development for a representative chondrichthyan species. By preserving whole embryos in three dimensions, micro-CT imaging enables developmental morphologies to be visualized at high resolution and in near-native anatomical context. Together with the recently published epaulette shark genome, this developmental atlas helps establish the Epaulette shark for comparative anatomical, developmental, and genomic studies.

## Introduction

Chondrichthyans, the most ancient surviving group of gnathostomes, are critical for insights into the evolution of the vertebrate body plan. Chondrichthyan reproduction and embryology have a long history of study, dating back to Aristotle (384-322 B.C.) who described oviparous versus viviparous reproductive strategies, egg case structure, and aspects of embryonic development (reviewed in Wourms and Demski 1993; Wourms 1997). In the nineteenth and early twentieth centuries, descriptive works characterizing the development and anatomy of elasmobranchs took a prominent role in the study the vertebrate body plan (e.g., Balfour, 1878; Dohrn 1884: Gegenbaur, 1865; Scammon, 1911; Goodrich 1958), laying the groundwork for modern molecular studies (e.g., Neyt et al, 2001; Dahn et al, 2007; Freitas et al, 2006; Gillis et al, 2009; Adachi et al, 2012; Onimaru et al, 2015; Nakamura et al, 2015; Gillis et al, 2016; Onimaru et al, 2016; Oshaughnessy et al, 2015; Barry et al, 2017; Tulenko et al, 2017; Sleight and Gillis, 2020; Onimaru et al, 2021; Hirschberger et al, 2021; Kusakabe et al, 2021; Kuroda et al, 2021; Rees et al, 2023). Efforts to sequence the genomes of several chondricthyan species (Pearce et al, 2021; Nishimura et al, 2022) have further advanced comparative and functional genomic approaches, providing insight into the evolution of features such as batoid fins (Marletaz et al 2023), Hox cluster organization (Hara et al 2018), the genetic systems that regulate neuronal circuits for locomotion (Yoo et al, 2022); and vertebrate germ line mutation rates (Sendell-Price et al 2023), as just a few examples. Despite these advances, studies of chondrichthyan development have been relatively hindered compared to established bony fish model systems (e.g., zebrafish, xenopus, chick, mouse) by lack of easy, regular access to materials.

To further facilitate the study of shark development, we propose the Epaulette shark (*Hemiscyllium ocellatum*) as a tractable laboratory model system. The Epaulette shark is a nocturnal, slender-bodied, oviparous shark that inhabits the coral reefs of the northeastern coast of Australia (Heupel and Bennett, 1998; Allen et al., 2016; Dudgeon et al., 2020). Epaulette sharks are relatively small, reaching a size of approximately 80–107 cm as adults, and are decorated with a variety of dark pigmentation spots, as well as a prominent “epaulette” just above their pectoral fins (**Figure 1A**; **Figure 6**). While Epaulette sharks have been of particular interest to researchers for decades due to their ability to ’walk’ along the sand and exposed reef flats (Goto, Nishida and Nakaya, 1999; Porter et al., 2022) and their tolerance of anoxic conditions (Chapman, Harahush and Renshaw, 2011; Routley, Nilsson and Renshaw, 2002; Wise, Mulvey and Renshaw, 1998), their development has come into recent focus (West and Carter, 1990; Payne, 2012; Johnson et al, 2016; Masselink et al, 2026; Wheeler et al, 2021; Thomas et al, 2023; Peele et al, 2026; Gaudi et al, 2026).

**Figure 1.**
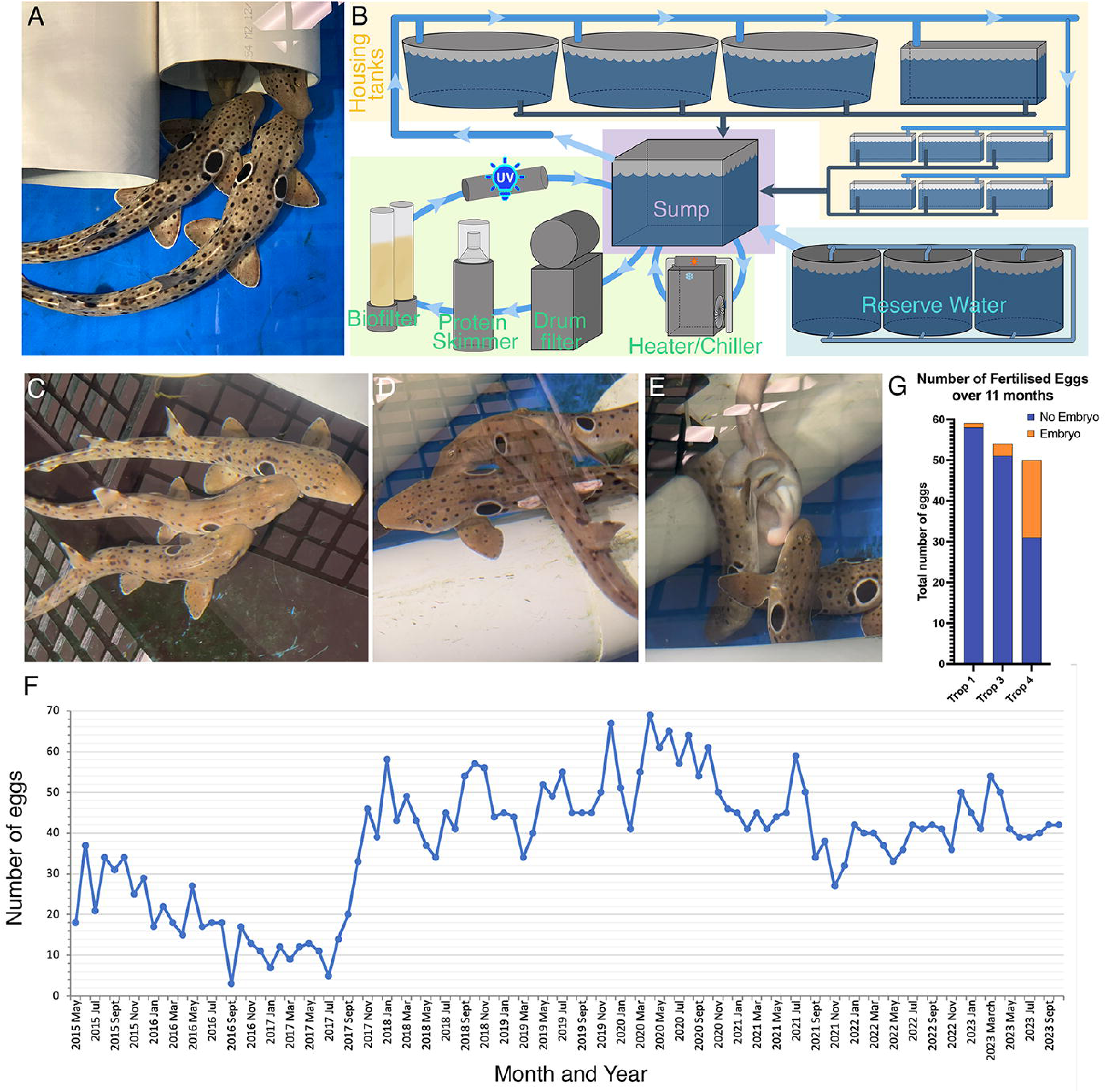
Brood colony of the Epaulette shark *Hemiscyllium ocellatum*. **A.** Example of adult epaulette sharks maintained as part of a captive brood stock. **B.** Schematic overview of marine system for maintaining adult sharks and eggs. Arrows indicate direction of water flow. Yellow box, adult and egg housing tanks; cyan box, reserve water tanks; purple box, sump; and green box, water filtration and processing. **C–E.** Time lapse of captive mating showing mating initiation in which the male bites the pectoral fin of the female (C), followed by rolling of the female (D), and insertion of the clasper (E). Note in panel (D), claspers can be seen extending away from the body. **G.** Frequency of successful embryo production varies with mating partners. Trop1, 2, and 3 refer to separate tanks containing unique sets of sharks. **E.** Egg production does not vary seasonally.

Detailed 3-D atlases of developmental morphologies are fundamental resources for developmental biology, providing context for studies of molecular patterning, tissue differentiation, and organogenesis. While 3-D anatomical atlases of development have been at least partially generated for human (Yamada et al, 2012; De Bakker et al, 2016), mouse (Baldock et al, 2003; Armit et al, 2017; Wong et al, 2012; Wong et al, 2015; Devine et al, 2022), avians (Ruffins et al, 2007; Wong et al 2013) and *Xenopus* (Laznovsky et al, 2024), and in some cases have been integrated with broader ontologies that include gene expression and function (e.g., emouse atlas, Armit et al, 2017), fewer resources exist for fish species. Serial sections and optical projection tomography have been used to produce 3-D atlases for zebrafish (Verbeek et al, 2000; Bryson-Richardson et al, 2007). Amoung cartilaginous fish, several general staging guides based on illustration or brightfield microscopy have been generated. These have included guides for the small-spotted catshark Scyliorhinus canilcula (Ballard 1993), Port Jackson shark *Heterodontus portusjacksoni* (Rodda and Seymour, 2008); winter skate Leucoraja ocellata (Maxwell et al, 2008), clearnose skate *Rostroraja eglanteria* (Luer et al, 2009); brownbanded bamboo shark *Chiloscyllium punctata* (Harahush et al, 2007; Onimaru et al, 2018), Atlantic sharpnose shark *Rhizoprionodon terraenovae* (Castro and Wourms, 1993), frilled shark *Chlamydoselachus anguineus* (Lopez-Romerao et al, 2020), Bonnethead *Sphyrna tiburo* (Byrum et al, 2023), and the Elephant fish *Callorhincus milii* (Didier et al 1998). General 3-D anatomical atlases of development for representative chondrichthyans, however, are lacking.

Here we provide resources for establishing Epaulette sharks as a tractable laboratory model system for studying shark development, including a description of our brood colony setup and husbandry, as well as a high resolution, micro-CT based three-dimensional atlas of post-neurula through hatching stages. These resources, together with the recently published Epaulette genome (Sendell-Price et al 2023), facilitate the use of Epaulette shark as a representative chondrichthyan in modern molecular studies of development.

## Results

### Adult Epaulette Shark Housing and Husbandry

#### Housing

A custom-built 22,000L closed marine system for maintaining a brood colony of adult epaulette sharks was established at Monash University that included three 5000L round tanks and one 2100L tank to house adult sharks; six 350L glass tanks to rear eggs; and a 2800L sump (**Figure 1A, B)**. Sea water was obtained by truck from Port Phillip Bay and supplemented with artificial sea water. Adults were housed at a maximum stocking density of ten sharks per tank. An overview of the filtration system and water path is presented in **Figure 1B**. In brief, seawater is the system empties from the adult and egg tanks (**Figure 1B**, yellow box) into a sump (**Figure 1B**, purple box), from which it then flows through a large drum filter (Fresh by Design) to remove particulate matter, a protein fractionator (RK2 Systems RK75PE) to remove organic waste, and a biofilter (RK2 Systems FSF Fluid Sand Filter), as well as a UV light sterilizer and ozone generator for processing (Figure 1B Green Box). A chiller (Toyesi) and heating unit (Elecro) were used to regulate temperature. Water parameters were monitored for temperature, pH, dissolved oxygen, salinity, nitrates, nitrites and ammonia (**Supplementary Table 1**) (see Mohan and Aiken, 2004 for detailed descriptions of chondrichthyan aquaculture). A 12-hour daylight cycle with graded dawn and dusk illumination was maintained using a series of LED lamps positioned above each tank and regulated by Casambi software. Large circular lids equipped with access hatches were used to cover each tank to prevent jumping. Because of their unique spot patterns, all sharks were photographed for identification cards.

#### Feeding

In the wild, epaulette sharks hunt for food in rock pools exposed during low tides, eating a diet of small fish, worms, crabs and other invertebrates (Heupel and Bennett, 1998). Given their broad diet, captive Epaulette adults were fed commercially available frozen bait fish, including pilchard, whiting, and glassies, as well as invertebrates such as pipis, mussels and squid. In total, sharks were fed four times per week, each receiving approximately 34 grams per feeding, which totals approximately 135g per week [or 5-7% of average Epaulette shark adult body mass (Wheeler et al., 2023)]. Feeds were supplemented with Elasmotabs (Vetafarms), a specialized dietary supplement for captive elasmobranchs.

#### Breeding Setup and Mating

Healthy adult Epaulette males have a relatively slim trunk, whereas females vary in girth throughout the reproductive cycle between egg laying (**Supplementary Figure 1**) (Wheeler et al., 2023). Sharks were housed with a mix of one to eight females and one to two males. Matings occurred spontaneously and did not require specific triggers. An example of captive Epaulette mating behaviour is illustrated in Figure 1C–E, and initiates with a male biting the female’s pectoral fin (**Figure 1C**), followed by rolling of the female (**Figure 1D**), and insertion of the clasper into the female cloaca (**Figure 1E**). The siphon sacs, which are paired structures that extend from the claspers along the ventral body wall, inflate with sea water during mating to propel sperm into the female (Gilbert 1972), prior to release of the female.

#### Egg Production and Fertilization Rates

To determine if egg production varied with season, total egg production was plotted against time **(Figure 1F)**. From these data, there are no clear indications of seasonal differences in egg laying. Fecundity, however, varied between isolated mating pairs (**Figure 1G**), suggesting reproductive quality may differ between individual sharks, or potentially mate choice may be important. While female mate selection has been proposed for nurse sharks and scalloped hammerheads, relatively little is known about mate choice in chondrichthyans.

### Three-dimensional Atlas of Epaulette Shark Developmental Morphologies

Detailed characterizations of embryonic anatomy across developmental timepoints provide an essential framework for molecular developmental studies of patterning and morphogenesis. Here we provide a three-dimensional anatomical staging atlas for Epaulette shark based on micro-computed tomography (micro-CT) scans complemented with brightfield microscopy. This staging guide is modified from the staging frameworks proposed by Ballard, Mellinger and Lechenault (1993b) for *Scyliorhinus canicula* (catshark) and Onimaru et al. (2018) for *Chiloscyllium punctatum* (bamboo shark). Micro-CT scans with iodine enhanced contrast to visualize soft tissues (Metscher, 2009) were generated for stages 18 (post-neurula) through 38 (hatching stage). Prior to Stage 18, embryos did not retain sufficient iodine for microCT scanning and were therefore not suitable for inclusion [but see *T. Sauka–Spengler, J.–L. Plouhinec, and S. Mazan 2004;* Kopsch 1950; Ballard 1993; Onimaru et al. (2018) for descriptions of chondrichthyan gastrulation]. The number of specimens examined for each developmental stage is summarized in **Supplementary Table 2**.

An overview of Epaulette stages is presented in **Figures 2** and **3**. Although the embryonic tail is normally curled between stages 18 and 24, it appears straightened in micro-CT scans for Stages 22–24 in Figure 2B as a mounting artifact because embryos were positioned to minimize scanning diameter and maximize resolution. Developmental stage showed a logarithmic relationship with days post-deposition (**Figure 4A**), whereas embryo length increased linearly over the same period (**Figure 4B**). Eggs maintained at 25-26 °C hatched after approximately 4.5–5 months of incubation.

**Figure 2.**
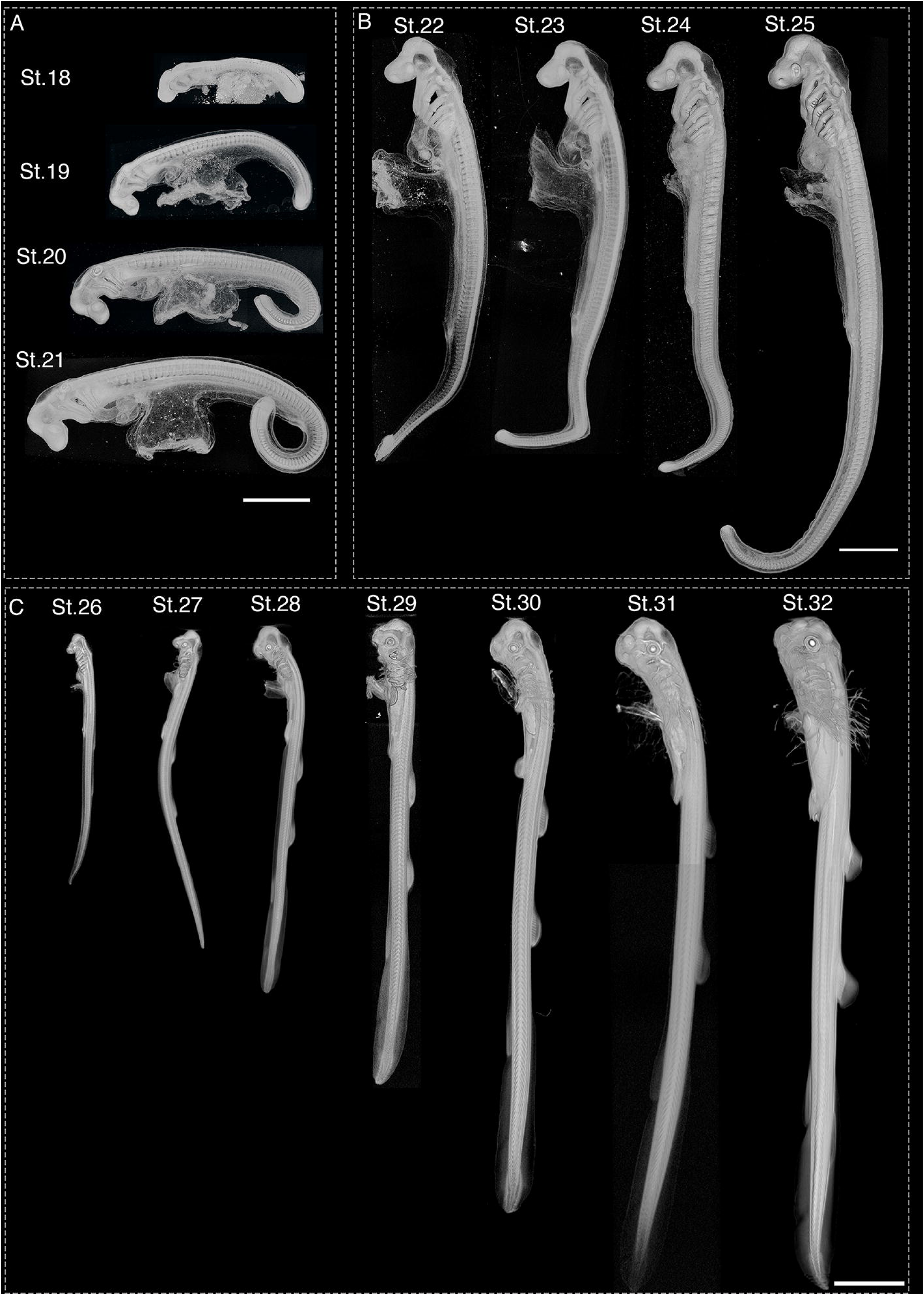
Overview of Epaulette shark developmental stages 18–32. MicroCT based renderings of whole Epaulette shark embryos in left lateral view. **A.** Stages 18–21. Anterior is left, dorsal is up. **B.** Stages 22–25. Anterior is up, dorsal is right. **C.** Stages 26–32. Anterior is up, dorsal is right. In (A) and (B), scale bar = 1000um. In (C), scale bar = 5000um.

**Figure 3.**
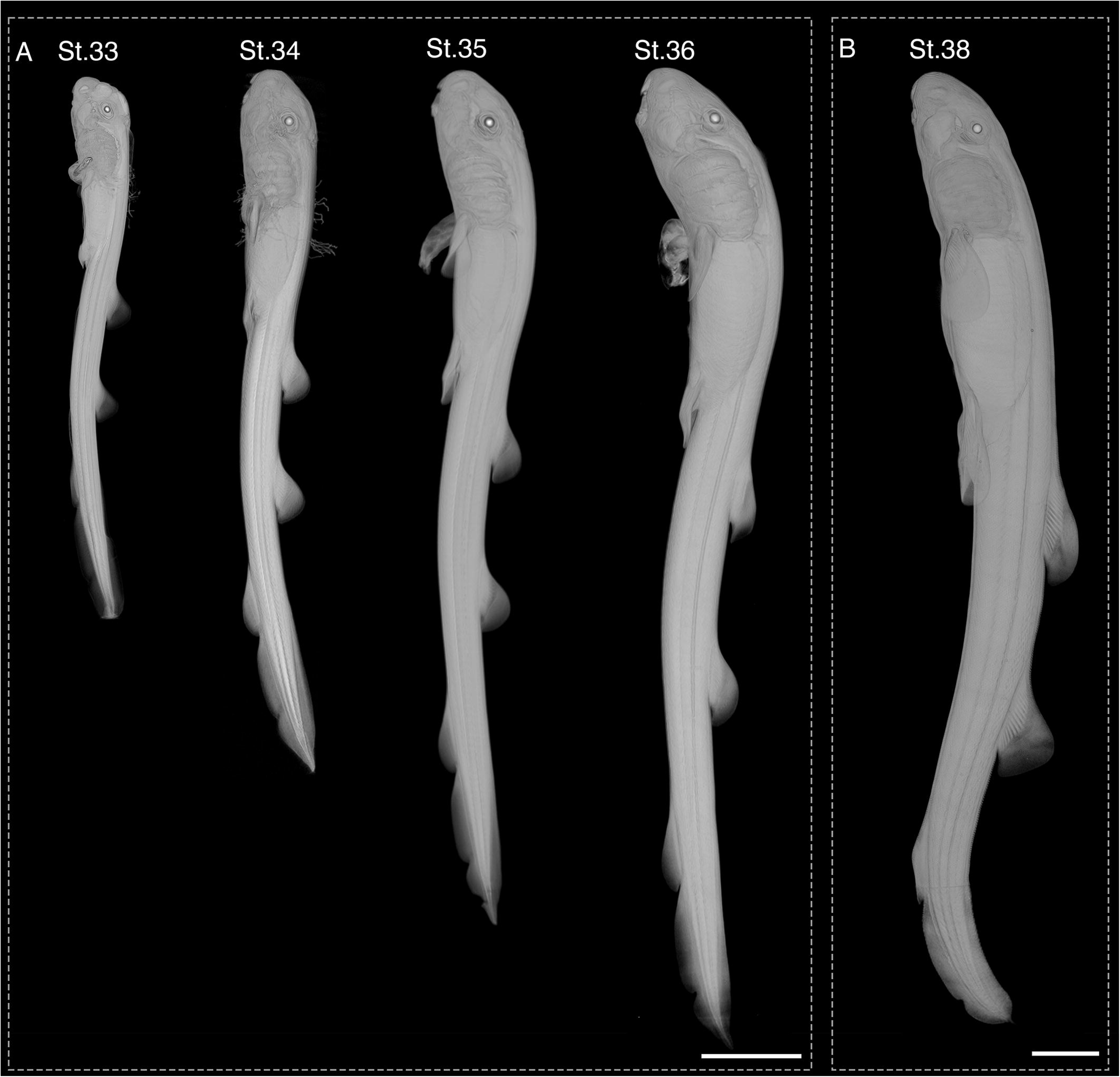
Overview of Epaulette shark developmental stages 33–38. MicroCT based renderings of whole Epaulette shark pre-hatching stages in left lateral view. For all images anterior is up, and dorsal is right. **A.** Stages 33–36. **B.** Stage 38, immediately prior to hatching. Scale bar = 10mm.

#### Stage 18

Stage 18 occurs at a mean of 10 days post deposition (dpd), with a range of 7–13 dpd (*n* = 5) (**Figures 2A, 4, and 5**). In lateral view, the forebrain, midbrain and hindbrain are visible, and a deep indentation is present in the hindbrain, corresponding to the position of rhombomere 3 (see Kuratani and Horigome, 2000). The optic vesicles are visible toward the anterior-most margin of head as outward swellings of the forebrain, though the width of the head remains relatively narrow. Rendered sections through the developing head show the internal shape of the early optic vesicles and forebrain **(Supplementary Figure 2).**

**Figure 4.**
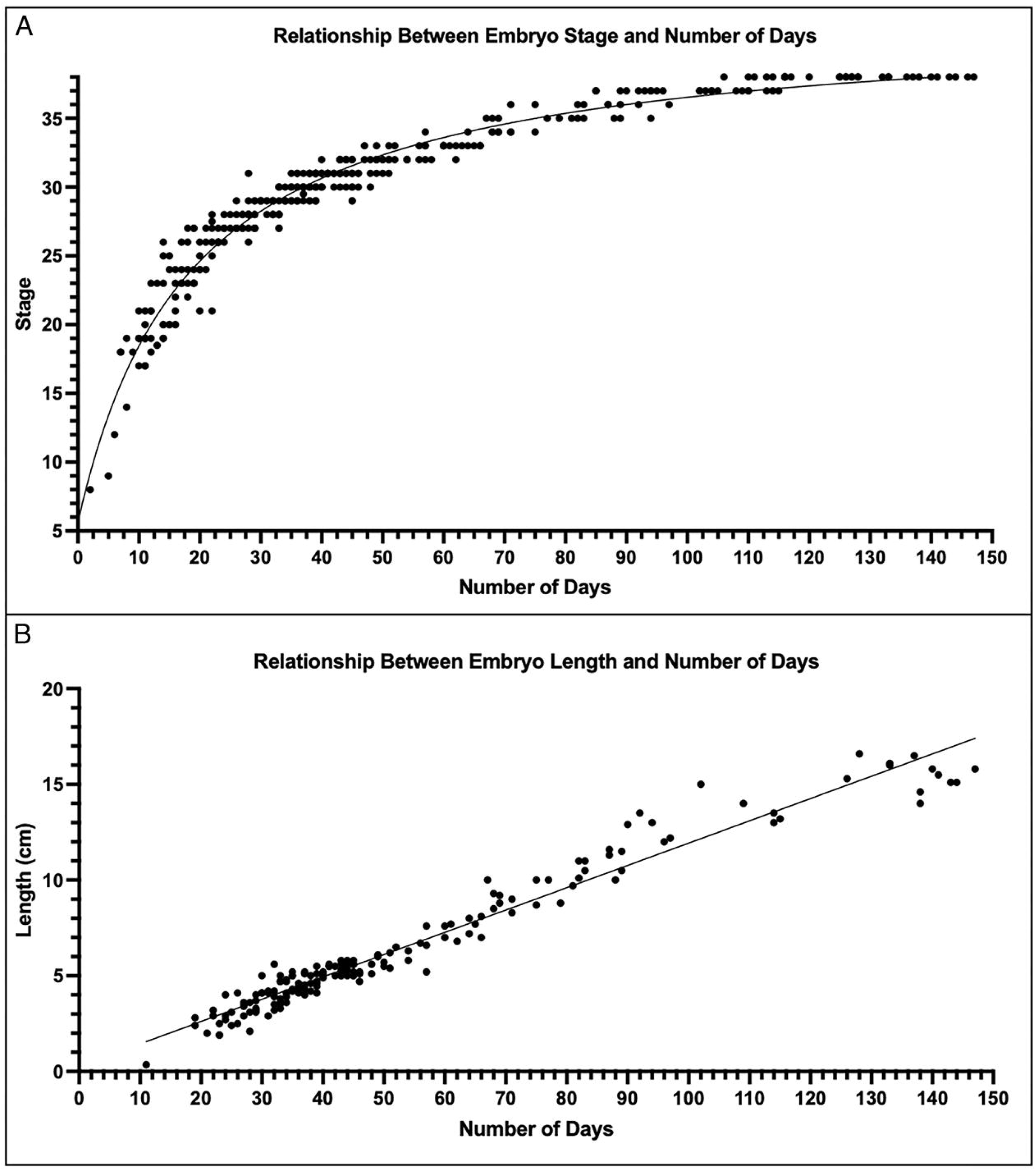
Relationship between developmental stage, embryo length, and age. **A.** Developmental stage shows a logarithmic relationship with days post-deposition. Non-linear regression: *R*² = 0.9558, *n* = 352. **B.** Embryo length increases linearly with age. Simple linear regression: *Y* = 0.1165*X* + 0.2779, *R*² = 0.9584, *n* = 174.

**Figure 5.**
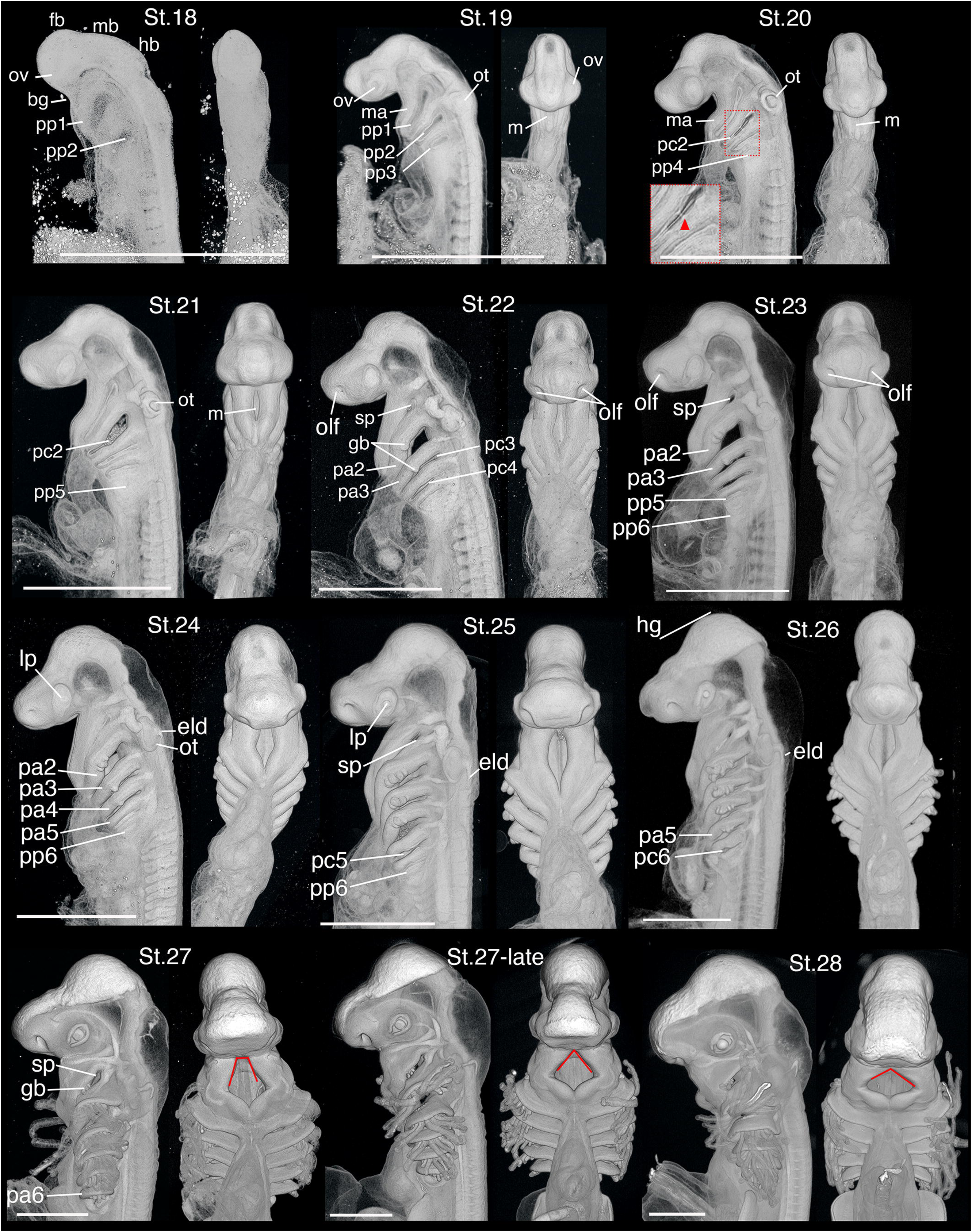
Development of the head at stages 18–28. For each stage, the head and anterior trunk are shown in left lateral (left panel) and ventral (right panel) views. Inset for stage 20 highlights discontinuous perforation (red arrowhead) of second pharyngeal cleft (pc2). In stages 27 and 28, red lines indicate mouth angle. bg, buccal groove; eld, endolymphatic duct; fb, forebrain; gb, gill bud; hb, hindbrain; lp, lens placode; m, mouth; ma, mandicular arch; mb, midbrain; ot, otic capsule; ov, optic vesicle; pa, pharyngeal arch 2; pa3, pharyngeal arch 3; pa4, pharyngeal arch 4; pa5; pharyngeal arch 5; pa6, pharyngeal arch 6; pc2, pharyngeal cleft 2; pc3, pharyngeal cleft 3; pc4, pharyngeal cleft 4; pc5, pharyngeal cleft 5; pc6, pharyngeal cleft 6; pp1, pharyngeal pouch 1; pp2, pharyngeal pouch 2; pp3, pharyngeal pouch 3; pp4, pharyngeal pouch 4; pp5, pharyngeal pouch 5; pp6, pharyngeal pouch 6; sp, spiracle.

**Figure 6.**
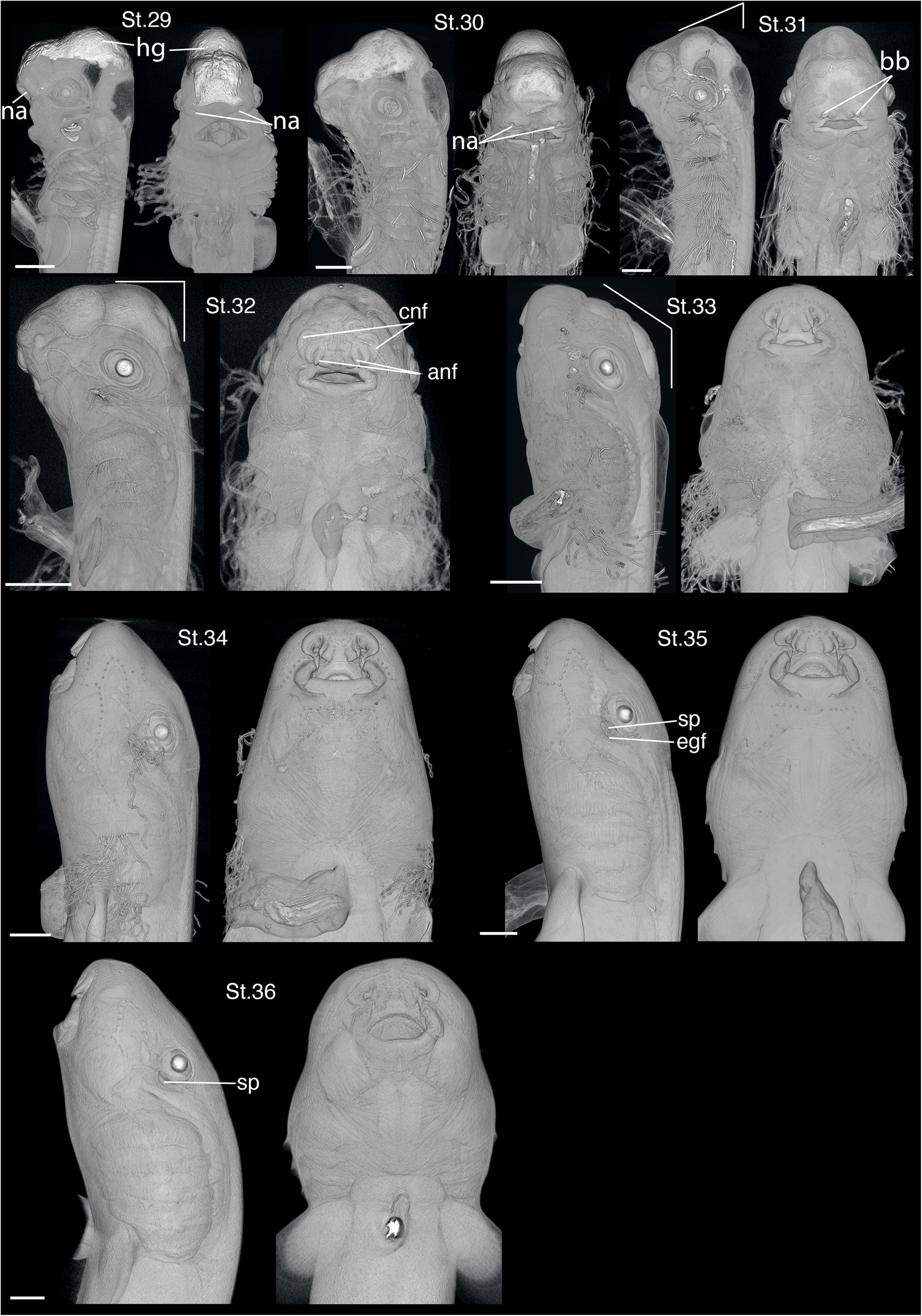
Development of the head at stages 29–35. For each stage, the head and anterior trunk are shown in left lateral (left panel) and ventral (right panel) views. For stages 29–31 the scale bar = 1mm. Angle that the front of the head makes with the body is indicated for stages 31 (acute), 32 (right angle) and 33 (obtuse angle). For stages 32–36 the scale bar = 2mm. anf, anterior nasal flap; bb, barbels; cnf, circumnarial folds; egf, external gill filaments; hg, hatching gland; na, nares; sp, spiracle.

Metamerization of the pharynx has initiated by this stage, as the first and second pharyngeal pouches form swellings that appear slightly translucent in lateral view. The buccal groove is visible as a shallow indentation between the optic vesicle and the first pharyngeal pouch.

Overall, the body has a broad base of attachment to the yolk (**Figure 2A; Supplementary Figure 2**). The heart tube is visible and appears straight. Approximately 27 pairs of somites extend along the trunk, ending in unsegmented paraxial mesoderm. The tail bud lifts off the yolk ball and curves slightly ventrally.

#### Stage 19

Stage 19 occurs at a mean of 11dpd, with a range of 8–14dpd (*n* = 10) (**Figures 2A, 4, and 5**). The optic vesicles have become more distinct outgrowths. The otic placodes are now visible as thickenings of the ectoderm on either side of the hindbrain and take the form of a slight invagination (Supplementary Figure 3). In contrast to the previous stage, the pharyngeal pouches now appear slit-like, and a third unopened pharyngeal pouch is visible. Additionally, the buccal groove forms a deep indentation with the anterior head, marking the position of the mouth.

There are approximately 49 pairs of somites along the length of the trunk. The heart tube is “S” shaped, forming the anlagen of the atrium and ventricle. The pronephros and kidney duct are visible as superficial structures adjacent to somites 4–11 (**Supplementary Figure 3**).

#### Stage 20

Stage 20 occurs at a mean of 14dpd, with a range of 11-16dpd (*n* = 7) (**Figures 2A, 4, and 5**). The invagination of each otic pit has deepened, forming an oval with well circumscribed margins (**Supplementary Figure 4**). The 4th pharyngeal pouch is now visible. Additionally, pharyngeal cleft 2 has opened from two discontinuous perforations. The mandibular arches frame the developing mouth, appears as a small slit that is visible in ventral view.

There are approximately 62 pairs of somites. The kidney duct extends further caudally relative to the previous stage. Interestingly, in scanned specimen Hoc#2052, the kidney duct lies adjacent to somites 7–16 on the left side and somites 7-18 on the right side (**Supplementary Figure 4**), indicating duct extension can occur asymmetrically during Epaulette shark development.

#### Stage 21

Stage 21 occurs at a mean of 15dpd, with a range of 10-22dpd (*n* = 7) (**Figures 2A, 4, and 5**). Each otic pit has partially constricted along its external margin, leaving an opening that is reduced in size relative to the forming capsule. The 5th pharyngeal pouch is now visible as a swelling. Pharyngeal cleft 2 opens broadly at its dorsal margin and narrows ventrally. By this stage the mouth is now open.

Approximately 72 pairs of somites extend along the trunk. The position of the cloaca is now clearly visible at the axial level of somites 35-41, but does not appear perforated (**Supplementary Figure 5**).

#### Stage 22

Stage 22 occurs at a mean of 17dpd, with a range of 16–18dpd (**Figures 2B, 4,** and **5**). The olfactory placodes are now visible in lateral and ventral views as shallow depressions along the anterior margin of the head (**Supplementary Figure 6**). The overlying ectoderm of the optic vesicles has begun to thicken and forms a lens placode that abuts the optic vesicle, which has a concave shape (**Supplementary Figure 6**). The otic pits have closed forming capsules that are continuous with a short endolymphatic duct that opens directly at the overlying ectoderm (**Supplementary Figure 6**). The 5^th^ pharyngeal pouch forms a slit-like shape, and pharyngeal clefts 2, 3, and 4 are open. A single gill bud is visible along the caudo-lateral margins of pharyngeal arches 2 and 3.

#### Stage 23

Stage 23 of development occurs at a mean of 16dpd, with a range of 12-19dpd (*n* = 10) (**Figures 2B, 4,** and **5**). The olfactory placode appears deeper relative to the previous stage. The 6^th^ pharyngeal pouch is visible as a swelling, and the first pharyngeal cleft, the spiracle, has opened, though the timing of this appears to be variable. Three small gill buds are now visible on pharyngeal arch 2. The mouth forms a long, narrow opening that when viewed ventrally appears roughly diamond shaped.

#### Stage 24

Stage 24 of development occurs at a mean of 18dpd, with a range of 15-21dpd (*n* = 8) (**Figures 2B, 4,** and **5**). The position of the lens is now clearly discernible in the centre of the eye. The olfactory placode continues to become a more pronounced pit. The endolymphatic duct appears to stretch posteriorly from the otic capsule (**Supplementary Figure 7**). The 6th pharyngeal pouch is more well defined, and a single gill bud is now present on pharyngeal arch 4.

The kidney duct extends along the trunk to the level of the cloaca, where it forms a “Y” shape, with one branch extending towards the cloaca and the other positioned adjacent to the somites just caudal to the cloaca (**Supplementary Figure 7**).

#### Stage 25

Stage 25 occurs at a mean of 18dpd, with a range of 14-22dpd (*n* = 4) (**Figures 2B, 4,** and **5**). The lens appears as a well circumscribed structure now that is distinct from the ectoderm **(Supplementary Figure 8)**. The 6th pharyngeal pouch appears slit-like, but remains closed. Both pharyngeal cleft 5 as well as the spiracle are open. Five gill buds are now present on pharyngeal arch two, and three gill buds are present on both the third and fourth pharyngeal arches. In ventral view, the mouth appears pentagonal in shape.

The lateral line is visible along the trunk, branching from the post-otic ganglion and extending to the axial level of somite 8 on the left side of the body **(Supplementary Figure 8)**. Similar to the kidney duct described for Stage 20, the lateral-line can also exhibit slight asymmetry across the midline, extending half a somite further on the right side of the body in specimen Hoc#1516. Distinct loops are visible in the gut tube, marking the onset of spiralling, as seen in both lateral view and rendered cross section (**Supplementary Figure 8**).

#### Stage 26

Stage 26 occurs at a mean of 21dpd, with a range of 14-28dpd (*n* = 13) **(Figures 2B, 4, and 5).** In lateral view, the endolymphatic duct appears “L” shaped as it extends from the otic capsule (**Figure 3**). Pharyngeal cleft six is now open. Seven gill buds are now present on pharyngeal arch two, five gill buds pharyngeal arch three, three gill buds on pharyngeal arch four, and a single gill bud on pharyngeal arch five.

The lateral line extends further caudally than the previous stages, reaching somite 20 (**Supplementary Figure 9**). The pectoral fins are now present as small outgrowths on either side of the ventral flank just caudal to the heart and above the yolk stalk (**Figure 7**). The morphology of the ventral margin of the somites has changed from the previous stage. The somites located above the pectoral fins have begun to form two distinct extensions, termed muscle buds, ventrally (**Supplementary Figure 9**). The pelvic fins are also visible as slight ridges on either side of the cloaca, though appear smaller than the pectoral fins at this stage (**Figure 7**). The first and second dorsal fins are discernible as regionalised thickenings of the median fin fold (**Figure 7**).

**Figure 7.**
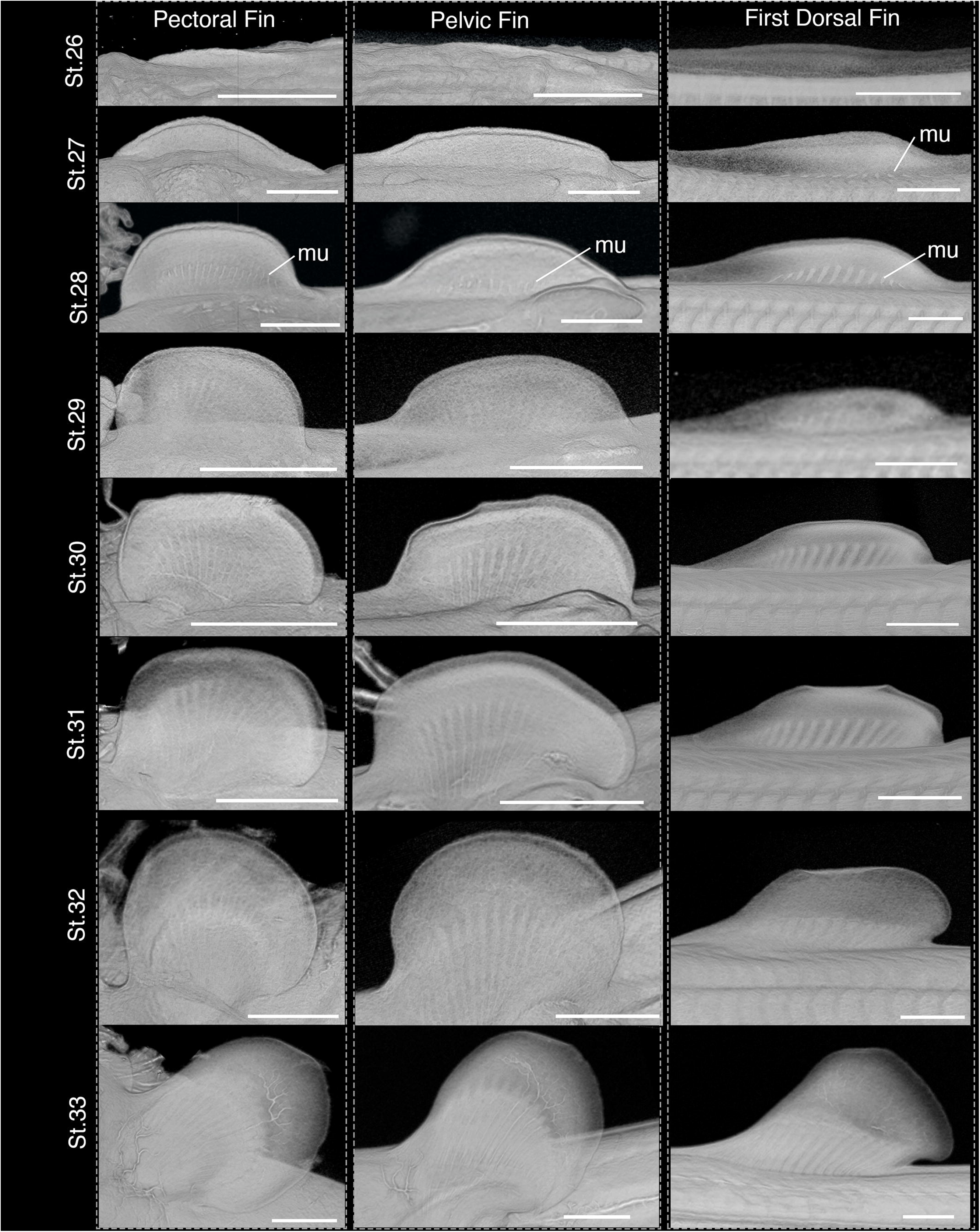
Development of the pectoral, pelvic and dorsal fins. Pectoral and pelvic fins are in ventral view. The first dorsal fins are in left lateral view. For all fins, anterior is left. mu, muscle buds.

#### Stage 27

Stage 27 occurs at a mean of 25dpd, with a range of 18–33dpd (*n* = 23) **(Figures 2B, 4, and 5).** The space between the maxillary processes has narrowed anteriorly, making the mouth appear pentagonal at early Stage 27 to square shaped by late Stage 27 in ventral view. The hatching gland is visible as an increased density along the anterior margin of the head (Ballard, Mellinger and Lechenault, 1993b). Gill buds are present on both the spiracle and pharyngeal arch 6, extending slightly past the margin of the first and sixth arches, respectively. The gill buds on arches 2–5 have increased in length, forming external gill filaments that extend away from the head.

The pectoral and pelvic fins are larger than the previous stage and appear round in ventral view (**Figure 7**). Somitic muscle buds extend to the proximal base of the paired fins early in stage 27 and into the fin by late Stage 27 **(Supplementary Figures 10 and 11**). Unlike the previous stage, the somites along the trunk between the pectoral and pelvic fins have also begun to form ventral paired projections (**Supplementary Figures 10 and 11**). The first and second dorsal fins are now well-defined outgrowths from the dorsal median fin fold, and the anal fin is visible as a thickening of the ventral median fin fold (**Supplementary Figure 10**). By late stage 27, the muscle bud projections extend from the dorsal margins of the somites into the dorsal fins. However, unlike the paired fins, each somite provides only a single myogenic projection to the dorsal fins (visible in lateral view and cross section) **(Supplementary Figure 11)**.

#### Stage 28

Stage 28 occurs at a mean of 29dpd, with a range of 22-33dpd (*n* = 24) **(Figures 2B, 4, and 5)**. The eye has gone from oval to a more rounded shape. Elongate external gill filaments exit the spiracle and pharyngeal cleft 6. The mouth appears antero-posteriorly compressed relative to the previous stage, now forming a diamond shape that is wider than long.

In ventral view, the posterior margin of the pectoral and pelvic fins form an obtuse angle with the body wall (**Figure 7**). Similarly, the posterior margins of the first and second dorsal fins form an obtuse angle with the trunk (Figure 7). The surrounding median fin fold has regressed relative to the previous stage, though remains present.

#### Stage 29

Stage 29 occurs at a mean of 34 dpd, with a range of 26-45 dpd (*n* = 39) (**Figures 2C, 4, and 6)**. Though not distinguishable by x-ray, the first pigmentation is visible in the eye by brightfield microscopy (**Figure 8**). The olfactory pits have deepened, forming distinct chambers (**Supplementary Figure 13**). A single large nare is present at the opening of each chamber.

**Figure 8.**
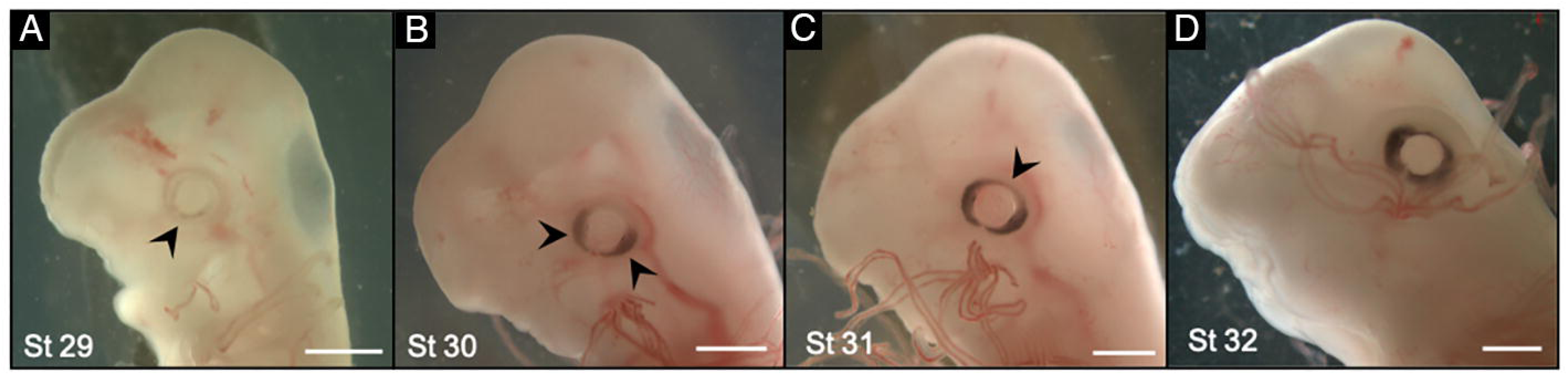
Formation of Eye Pigmentation. **A-D.** Pigmentation first appears in the eye at stage 29 (A) and extends along either side of the lens by stage 30 (B) before encircling the lens by stage 31 (C, arrowhead indicates dorsal closure of pigment ring). The pigment continues to surround the lens and forms a thicker circle from stage 32 (D) onwards. Scale bar 1mm.

In ventral view, the pectoral fins create a right angle with the body (**Figure 7**). Muscle buds now extend into the first dorsal fin. The anal fin is visible as a regional thickening of the ventral median fin fold, and at the very caudal tip of the tail the primordia of dermal denticles are visible (**Supplementary Figure 13**).

#### Stage 30

Stage 30 occurs at a mean of 39dpd, with a range of 33-48dpd (*n* = 31) (**Figures 2C, 4, and 6)**. By stage 30, pigment extends along the anterior and posterior margins of the eye but does not yet form a complete circle (**Figure 8**). The external nares constrict in the middle but remain open at the medial and lateral margins (**Supplementary Figure 14**). At this stage, the olfactory rosettes become distinct as a series of parallel grooves visible along the posterior epithelium of the olfactory chamber (**Supplementary Figure 14**). Flattening of the lower jaw now makes the mouth appear more compressed relative to the previous stage.

The posterior margin of the pectoral fin forms an acute angle with the trunk (**Figure 7**). The posterior margin of each dorsal fin forms a slightly obtuse angle with the tail. An indentation marks the posterior margin of the anal fin, though it remains continuous with the median fin fold anteriorly and caudal fin fold posteriorly (**Supplementary Figure 14**).

#### Stage 31

Stage 31 occurs at a mean of 41dpd, with a range of 28-51dpd (*n* = 35) **(Figures 2C, 4, and 6)**. The anterior margin of the head makes an acute angle with the axis of the body. Pigment in the eye now forms a complete circle around the lens **(Figure 8)**. The constriction along the centre of the nares is more pronounced, and the olfactory chambers form folds **(Supplementary Figure 15**). Barbels are now visible as ventral projections just anterior to the nares.

The posterior margin of the first dorsal fin now forms an acute angle with the tail (**Figure 7**). A deep indentation marks the posterior margin of the anal fin, which remains continuous with the fin-fold posteriorly (**Supplementary Figure 15**). In males, the claspers are first visible at stage 31 as swellings at the postero-medial margin of the pelvic fins (**Figure 9**).

**Figure 9.**
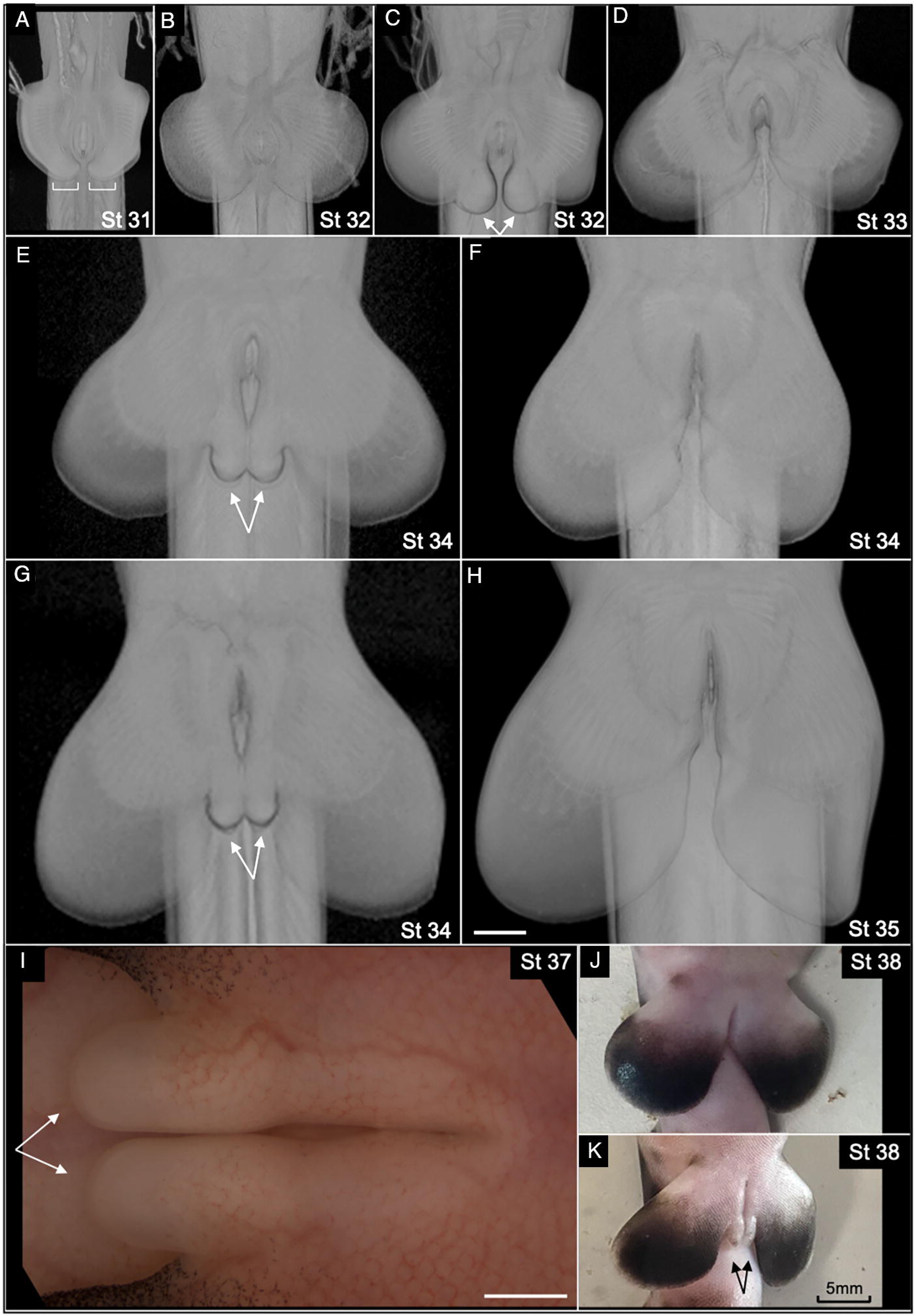
Clasper Development. **A–H.** MicroCT scans of stage 32-35 pelvic fins in ventral view. Anterior is up in all panels. **I–K.** Brightfield images of stage 37 (I, high magnification) and stage 38 (J,K) pelvic fins are shown in ventral view. In panel (I), anterior is to the right. In (J) and (K), anterior is up. Male pelvic fins are shown in A, C, E, G, I, and K. Female pelvic fins are shown at similar stages for comparison in B, D, F, H, and J. The developing claspers are first visible at stage 31 in males as swellings along the posterior edge of the pelvic fins (A, brackets). By stage 32 the claspers are distinct rounded protrusions (C, arrows), which become more elongate at later stages (E, G, I, J), but did not extend past the posterior margins of the pelvic fins. Scale bars, 1mm (A–I), 5mm (J, K).

#### Stage 32

Stage 32 occurs at a mean of 49dpd, with a range of 40-62dpd (*n* = 27) **(Figures 2C, 4, and 6)**. The anterior margin of the head now makes an approximate right angle with the axis of the body. The circumnarial fold and anterior nasal flaps are prominent around each barbel.

A deep indentation defines the lateral margin of each clasper, which are now more distinct protrusions that extend caudally from the pelvic fins (**Figure 9**).

#### Stage 33

Stage 33 occurs at a mean of 59dpd, with a range of 47-66 dpd (*n* = 17) **(Figures 3, 4, and 6)**. The anterior margin of the head now makes an obtuse angle with the axis of the body. The circumnarial fold, barbels and anterior nasal flap are more pronounced.

The central axis of the pectoral fin is directed postero-laterally (**Figure 7**). In addition to morphological features, from stage 33 onwards pigmentation patterns on the body provide key features for identifying stage. Pigment begins to form on the posterior edge of the dorsal fins and behind the caudal margin of the eye (**Figure 10**).

**Figure 10.**
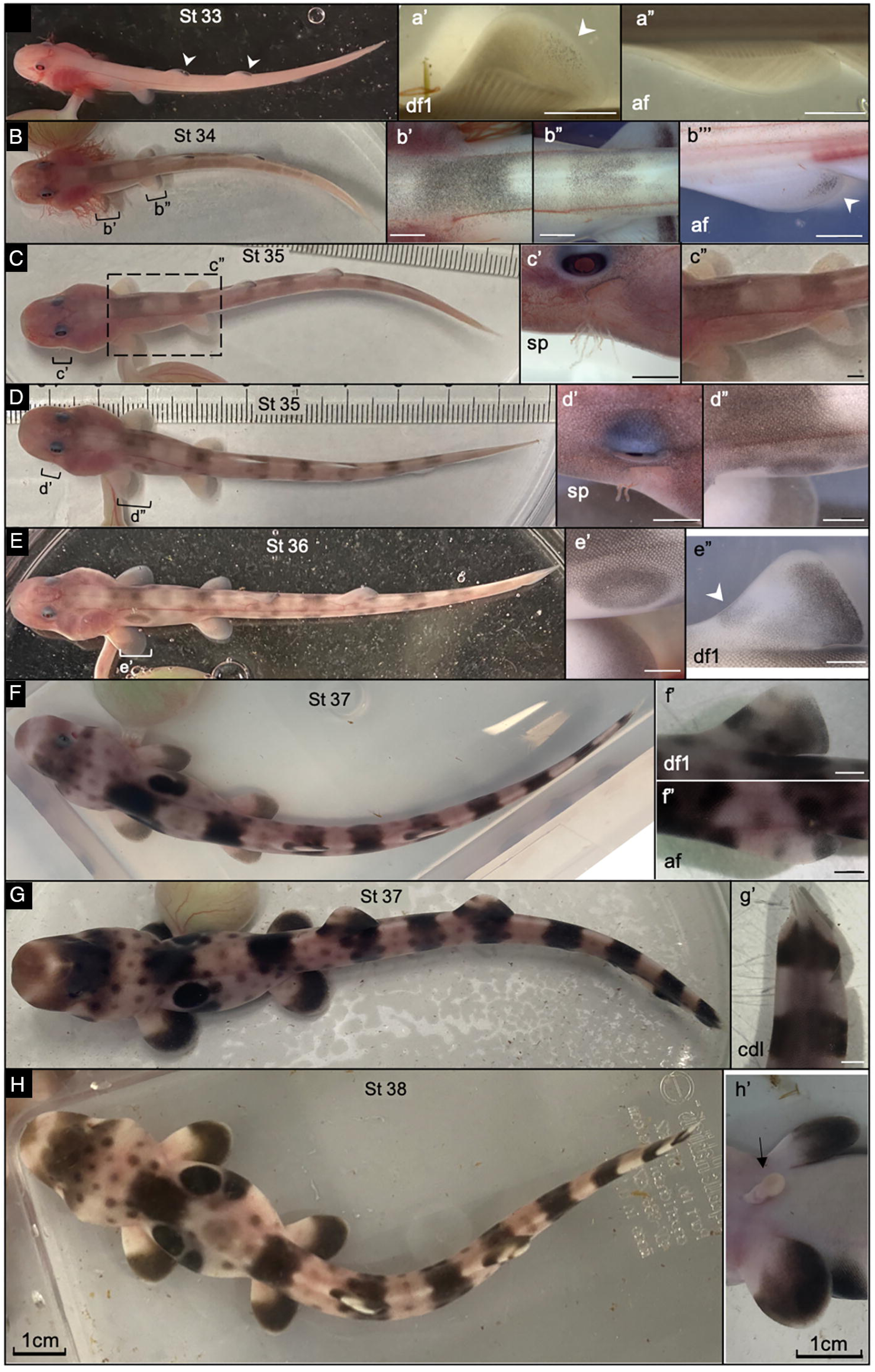
Development of body pigmentation. **A–H**. Dorsal view of entire pre-hatching stage epaulette shark removed from the egg case. Enlarged regions are indicated with brackets and are shown in dorsal (b’-e’, c”-d”), left lateral (a’,f’-g’, a”, e”-f”, b”’) and ventral (h’) views. The first fin pigmentation appears on posterior edge of the dorsal fins (df) at stage 33 (A, a’, arrowhead), but not on the anal fin (af) (a”). Bands of pigmentation appear along the trunk and tail by stage 34, along with the posterior edges of the pectoral, pelvic and anal fins (B, b”’, arrowhead). At Stage 35 external gill filaments remain present in the spiracle (sp), though appear reduced by late Stage 35 (c’,d’). The epaulette spot appears by late stage 35 (compare C and D). Additionally, feint spotting appears across the body at Stage 35 (D), which becomes more distinct by stage 36 (E) and a band of pigment begins to show on the anterior dorsal fin (e”, arrowhead). The pigment becomes darker and more defined at stage 37 (F and G). A band of pigment is visible at the tip of the snout, and defined bands continue to the end of the caudal fin (cdl) (g’). By Stage 38 the embryo has reabsorbed most of the yolk, noted by the rounded abdomen and small yolk stalk (h’, arrow). The pigment pattern remains similar to stage 37 with dark bands and spots visible across the entire body (H). Scale bars, 1cm for A-H and h’. Scale bars are 1mm for a’-g’, a”-f”, b”’.

#### Stage 34

Stage 34 occurs at a mean of 68dpd, with a range of 57-75dpd (n=9) **(Figures 3, 4, and 6)**. Long external gill filaments still extend outwards from the spiracle and gill clefts. Growth of the pelvic fins outpaces that of the claspers, such that the posterior margins of the fins extend further caudally than the claspers (**Figures 7, 9**). Pigment is now present along the postero-distal margin of the paired fins and becomes reduced proximal to the position of the apical fin-fold **(Figure 10)**. Feint bands are also beginning to form along the length of the trunk (**Figure 10**).

#### Stage 35

Stage 35 occurs at a mean of 80dpd, with a range of 67-94 dpd (*n* = 11) **(Figures 3, 4, and 6)**. External gill filaments remain present in the spiracle, but are largely reduced from the posterior gill slits. Spotting has begun to develop across the body by late stage 35, and by late stage 35 a distinct epaulette spot is present on the flank above the pectoral fins (**Figure 10**).

#### Stage 36

Stage 36 occurs at a mean of 85dpd, with a range of 71-97 dpd (*n* = 9) **(Figures 3, 4, and 6)**. The external gill filaments are lost from the spiracle. Spotting has become more defined and a band of pigment begins to develop on the leading edge of the dorsal fin (**Figure 10**). Additionally, the epaulette spot is now well circumscribed **(Figure 10)**.

#### Stage 37

Stage 37 occurs at a mean of 101dpd, with a range of 85-115dpd (*n* = 23) **(Figures 3, 4, and 6)**. The yolk ball is reduced in size. The pigmentation bands and spots becomes darker and more defined, with a band of colour developing on the tip of the snout (**Figure 10**).

#### Stage 38

Stage 38 occurs at a mean of 127dpd, with a range of 106-147dpd (*n* = 31) **(Figures 3, 4, and 6)**. This is the final stage prior to hatching, and is marked by complete resorption of the external yolk ball, retaining only the yolk stalk (**Figure 10)**. The abdomen appears rounded and contains yolk **(Figure 10)**.

### MicroCT Utility and special considerations

#### Developmental Morphologies of Somitic muscle buds during vertebrate appendicular muscle formation

High resolution MicroCT allows visualization of developmental morphologies in three dimensions that can be difficult to discern by wholemount brightfield microscopy alone. For example, the scans presented here highlight how the morphologies of the ventral somitic margins vary both regionally and over time, which has implications for understanding the evolution of vertebrate fin muscle patterning systems (Neyt et al; Cole et al; Tulenko et al 2013; Okamoto et al 2013; Turner et al 2019). During development, myogenic precursor cells originating from the hypaxial region of the fin-level somites populate the fin/limb where they differentiate into myofibres. In Stage 26 Epaulette sharks (a stage in which the paired fins are first visible as small ridges along the trunk), the ventral margins of the somites begins to bifurcate at the pectoral fin level (**Supplementary Figure 9**). By stage 27, these bifurcations take the shape of thin muscle buds, which also form further caudally along the inter-fin trunk and pelvic fin, but are absent caudal to the cloaca (**Supplementary Figure 10, 11)**. By Stage 28, muscle buds are visible extending well into the paired fins (See **Supplementary Figures 12**).

The epaxial somitic margins also contribute muscle buds to the dorsal fins, however, these occur as single, rather than paired, somitic projections (**Supplementary Figure 9–12**). Although the somites that form epaxial muscle buds are not restricted to the levels of the dorsal fins, they are absent from the anterior trunk, suggesting regionalized competency for their formation. Additionally, the directionality of epaxial muscle buds varies with axial level (**Figure 11**). All muscles buds from the anterior edge of the first dorsal fin to the anterior edge of the second dorsal fin (including inter-fin axial levels) are directed towards the first dorsal fin. Muscle bud orientation then abruptly shifts at the anterior margin of the second dorsal fin such that all buds become directed towards the second dorsal fin. Together, these observations suggest a system in which median fin signalling centres may direct cell behavior along the epaxial somites.

**Figure 11.**
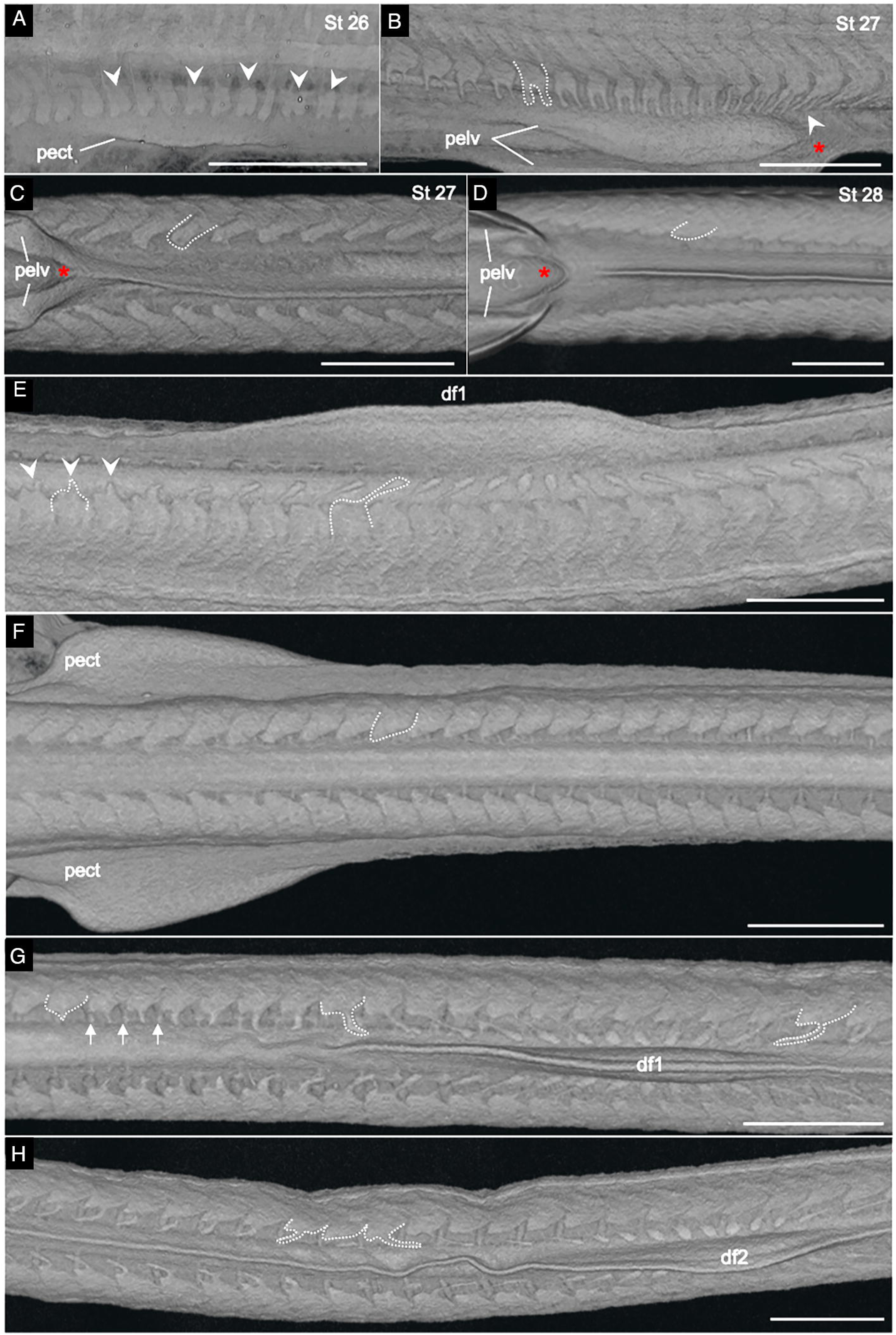
Developmental morphologies of somitic margins during appendicular muscle formation. MicroCT scans show morphologies of hypaxial or epaxial somitic margins at Stage 26 (**A**), stage 27 (**B, C, E–H**) and stage 28 (**D**). **A,B.** Left lateral view of the hypaxial somites at the pectoral (pect) (a) and pelvic (pelv) fin levels (B). Anterior is left. In **A**, ventral somite (arrowheads) margins begin to bifurcate coincident with when the paired fins first become visible. In Stage 27 (**B**), distinct paired muscle buds (highlighted with a white dashed line) are visible along the hypaxial somites of the trunk up to the cloaca (arrowhead). **C, D.** Ventral view of the tail. In Stage 27 (**C**) and Stage 28 (**D**), the somites posterior to the cloaca (red star) have a smooth ventral margin (white dashed line) and no paired muscle buds. Anterior is left. **E–H.** Left lateral (**E**) and dorsal **(F-H)** views of the dorsal, epaxial somite margins along the trunk. Anterior is left. In contrast to paired fins, a single muscle bud (white dashed line) extends from the dorsal epaxial margin of the somite towards dorsal fin 1 (**E,G**) or 2 (**H**) (df1, df2). While small muscle buds (arrowheads) are also visible just anterior to the first dorsal fin (**E**), they are not present further anteriorly (**F** shows trunk somites up to the level of the pectoral fins). **G, H.** Notably, muscle buds are visible along the dorsal margins of somites positioned in the interfin regions between df1 and df2, and show an abrupt change in directionality at the anterior margin of df2 (white dashed lines in **H** highlight epaxial muscle buds directed anteriorly versus posteriorly).

## Discussion

Several features of epaulette sharks make them well suited as a laboratory chondrichthyan model system. Their relatively small size and docile behaviour permit tractable aquarium husbandry. In captivity, epaulette sharks lay eggs throughout the year, with each female typically depositing two to four eggs per month (Payne, 2012; West and Carter, 1990). Because eggs are deposited at approximately the blastodisc stage, most stages of development can be accessed without sacrificing the mother. The 3-D anatomical atlas presented here, generated from micro-CT and brightfield datasets, provides a comprehensive overview of epaulette shark embryonic morphology. Together with the recently published epaulette shark genome assembly (Sendell-Price et al., 2023), this first detailed staging guide for the species establishes key resources for studies of epaulette shark development.

The developmental morphologies visualized by micro-CT provide new insights into shark development and can be used in comparative studies to better understand the evolution of the vertebrate body plan. For example, one of the prominent hypotheses for the evolutionary origin of paired fins is the lateral fin fold hypothesis, originally put forward in the nineteenth century (Balfour, 1881; Mivart, 1879; Thacher, 1877). According to this hypothesis, the paired pectoral and pelvic fins of jawed vertebrates arose from ancestral paired fin folds that extended along the length of the lateral trunk. Although this model has been widely argued in the literature, direct fossil evidence for a lateral fin fold remains contentious (Bemis and Grande, 1999; Coates, 1994; Coates, 2003). Historically, indirect evidence from comparative embryology has also been used to evaluate this hypothesis. Early descriptive work by Balfour (1878) suggested that the pectoral and pelvic fins of sharks and skates develop from a thickened ectodermal ridge extending along the trunk, and that this ridge was an embryonic remnant of an ancestral lateral fin fold. However, a more recent SEM study in the lesser spotted catshark, a shark belonging to the order Carcharhiniformes, did not observe the ectodermal thickening originally described (Tanaka et al., 2002). Given these conflicting observations, the epaulette shark, provides an additional phylogenetic data point for the order Orectolobiformes to evaluate this hypothesis. Similar to the catshark, there was no visible ectodermal thickening along the flank of epaulette shark, as the first signs of paired fin outgrowth were distinct localised ridges at pectoral and pelvic fin levels.

In addition to fin outgrowth along the body wall, the developmental morphologies of somites at fin and inter-fin levels have been discussed in the context of early paired appendage evolution (Goodrich 1930; Neyt et al, 2000). Moreover, current molecular models support model in which fin patterning systems first evolved in midline fins, and were then secondarily redeployed to the flank with the origin of paired fins. In tetrapods, myogenic precursor cells from limb-level somites migrate into the limb bud as de-epithelialized mesenchyme, where they differentiate to form the dorsal and ventral muscle masses (Deries and Thorsteinsdóttir, 2016). A similar process operates for pectoral fin muscle formation in zebrafish, a teleost (Haines and Currie, 2001; Neyt et al., 2000), suggesting a deep conservation with tetrapods. However, a different dynamic has been described for chondrichthyans. Early studies of the lesser spotted catshark describe that at the pectoral and pelvic fin level, epithelial muscle buds extend from the somites into the nascent fins to produce muscle (Cole et al., 2011; Goodrich, 1930; Neyt et al., 2000). During this process, the ventral somitic cells entering the fin express molecular markers of appendage muscle precursor cells, including the transcription factors Pax3 and Lbx1 (Kusakabe et al., 2020; Okamoto et al., 2017; Turner et al., 2019). Interestingly, it has also been proposed that shark muscle buds may undergo a transient de-epithelialization as they extend from the ventral somites (Okamoto et al., 2017). The microCT scans of epaulette sharks provides a visualisation of the gross morphologies of the somite margins across the different stages of fin development. In epaulette sharks, we observed that prior to Stage 26, the ventral margins of the somites appear rounded and smooth in both the trunk and tail. In contrast, as the paired fins form, the ventral margins of the hypaxial somites adjacent to the fins begin to bifurcate, giving rise to two muscle buds, which by early stage 27 can be seen as continuous extensions from the somites into the paired fins. Interestingly, these paired muscle buds also form along the inter-fin trunk in Epaulette sharks, but not in the tail. In vertebrates, the lateral plate mesoderm extends from the head-trunk interface to the cloaca (Durland et al., 2008; Prummel et al., 2019), and during development, the hypaxial margin of trunk level somites abut the lateral plate mesoderm. In Epaulette shark embryos, the presence of paired muscle buds only up to the cloaca suggests that lateral plate derived signals may have a role in regulating somitic bud formation. In amniotes, migratory somitic cell specification is influenced by both an intrinsic somitic Hox code as well as cues from the forming limb bud (Alvares et al., 2003). However in sharks, the potential signalling interactions between somitic and lateral plate mesoderm remain to be elucidated.

Our scans also reveal that while two buds extend from the ventral somite into paired fins, only a single bud appears to extend from the dorsal margin into the unpaired dorsal fins. How this relates to the early evolution of paired fin musculature remains an open question. Additionally, axial differences in dorsal muscle bud orientation between the inter-fin tail and the second dorsal fin suggest an abrupt patterning boundary at the anterior margin of the second dorsal fin. Furthermore, because dorsal somitic buds arise in the region between the first and second dorsal fin, it is clear the systems responsible for their induction are not limited to the fin regions exclusively.

Overall, the resources described here help establish the epaulette shark, *Hemiscyllium ocellatum*, as a valuable emerging model for investigating chondrichthyan development. More broadly, these data create a foundation for future comparative studies of developmental patterning, anatomical evolution, and the origins of key vertebrate traits from a phylogenetically informative cartilaginous fish lineage.

## Materials and Methods

### Microcomputed tomography and visualization

Prior to scanning, each sample was dehydrated and stained to enhance contrast with Ethanol-Iodine as previously described (Metscher, 2009). After staining, specimens were rehydrated and mounted in sealed plastic vessels for scanning. All scans were performed using a Zeiss Xradia 520 Versa system.

Sample visualisation, analysis, and image generation were performed using Avizo Software. Staging criteria are based on features and frameworks identified in the lesser spotted catshark (Scyliorhinus canicula) (Ballard, Mellinger and Lechenault, 1993b) and the brown banded bamboo shark (Chiloscyllium punctatum) (Onimaru et al., 2018).

