## Supplementary Tables and Figures for "3-D Ontogenetic Staging Atlas of the Epaulette Shark *Hemiscyllium ocellatum,* a Laboratory Model for Shark Development"

**Table 5.1.:** Water Quality Parameters

| Parameter | Range |
|---|---|
| Salinity | 30-35 ppt |
| pH | 8.0-8.4 pH |
| Dissolved Oxygen | 7-9 mg/L |
| Ammonia | 0 mg/L |
| Nitrite | <0.1 mg/L |
| Nitrate | <100 mg/L |
| Temperature | 24-26 °C |

**Supplementary Table 1. Epaulette shark marine system water parameters.**

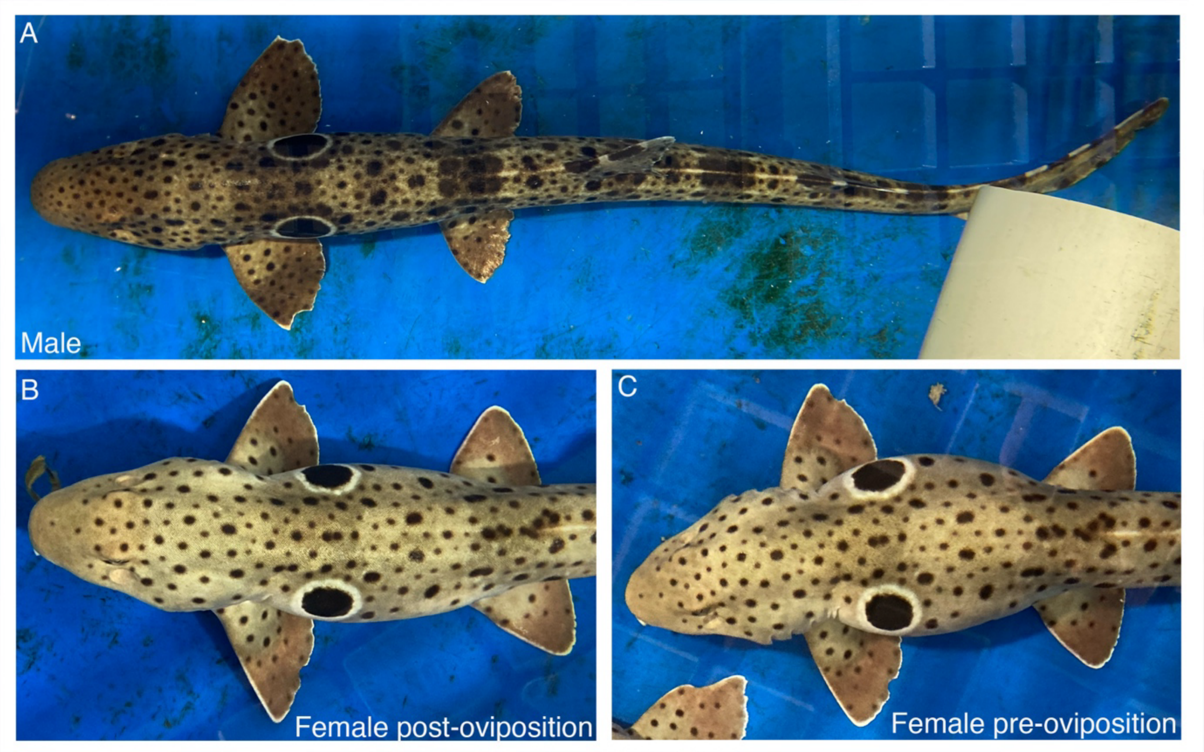


**Supplementary Figure 1. Male and female adult epaulette shark general body condition. A.** Typical adult male. **B, C.** Typical adult female at two different time points in reproductive cycle. Females have a relatively slim inter-fin trunk post-oviposition (B) and a wider girth during the egg encapsulation (C).

**Table 5.3.:** Epaulette Shark Development: Stage and Number of Days

| Stage | Mean Number of Days | Range of Days | Number of Samples |
|---|---|---|---|
| 8 | NA | 2 | 1 |
| 9 | NA | 5 | 1 |
| 12 | NA | 6 | 1 |
| 14 | NA | 8 | 1 |
| 17 | 11 | 10 - 11 | 3 |
| 18 | 10 | 7 - 13 | 5 |
| 19 | 11 | 8 - 14 | 10 |
| 20 | 14 | 11 - 16 | 7 |
| 21 | 15 | 10 - 22 | 7 |
| 22 | 17 | 16 - 18 | 2 |
| 23 | 16 | 12 - 19 | 10 |
| 24 | 18 | 15 - 21 | 8 |
| 25 | 18 | 14 - 22 | 4 |
| 26 | 21 | 14 - 28 | 13 |
| 27 | 25 | 18 - 33 | 23 |
| 28 | 29 | 22 - 33 | 24 |
| 29 | 34 | 26 - 45 | 39 |
| 30 | 39 | 33 - 48 | 31 |
| 31 | 41 | 28 - 51 | 35 |
| 32 | 49 | 40 - 62 | 27 |
| 33 | 59 | 47 - 66 | 17 |
| 34 | 68 | 57 - 75 | 9 |
| 35 | 80 | 67 - 94 | 11 |
| 36 | 85 | 71 - 97 | 9 |
| 37 | 101 | 85 - 115 | 23 |
| 38 | 127 | 106 - 147 | 31 |

**Supplementary Table 2. Number of eggs for each developmental stage**

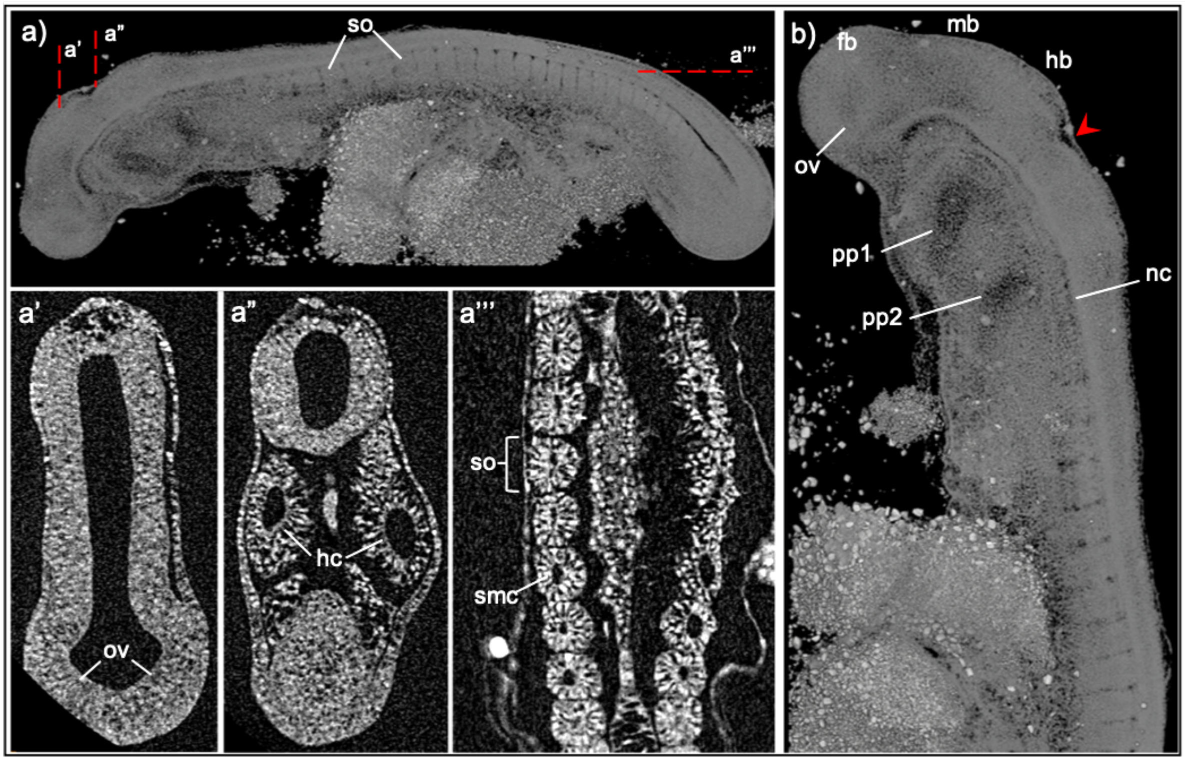


**Supplementary Figure 2. MicroCT of Stage 18 Epaulette Shark Embryo.** a. Left lateral view of a stage18 epaulette shark embryo in whole mount, head to the left tail bud to the right, showing the somites (so) along the trunk. a’-a”. Rendered cross sections through the head shows a bulging in the anterior fore brain where the optic vesicle (ov) will form (a’), and two head cavities (hc) (a”). Coronal section through the somites shows a distinct somitocoele at the center of the somites (a”’). b. Left lateral view of the head with visible fore (fb), mid (mb) and hind brain (hb), a distinct indent which aligns with the location of rhombomere 3 (red arrow), pharyngeal pouch 1 and 2 (pp1, pp2), optic vesicle and notochord (nc). Scale bar 500μm.

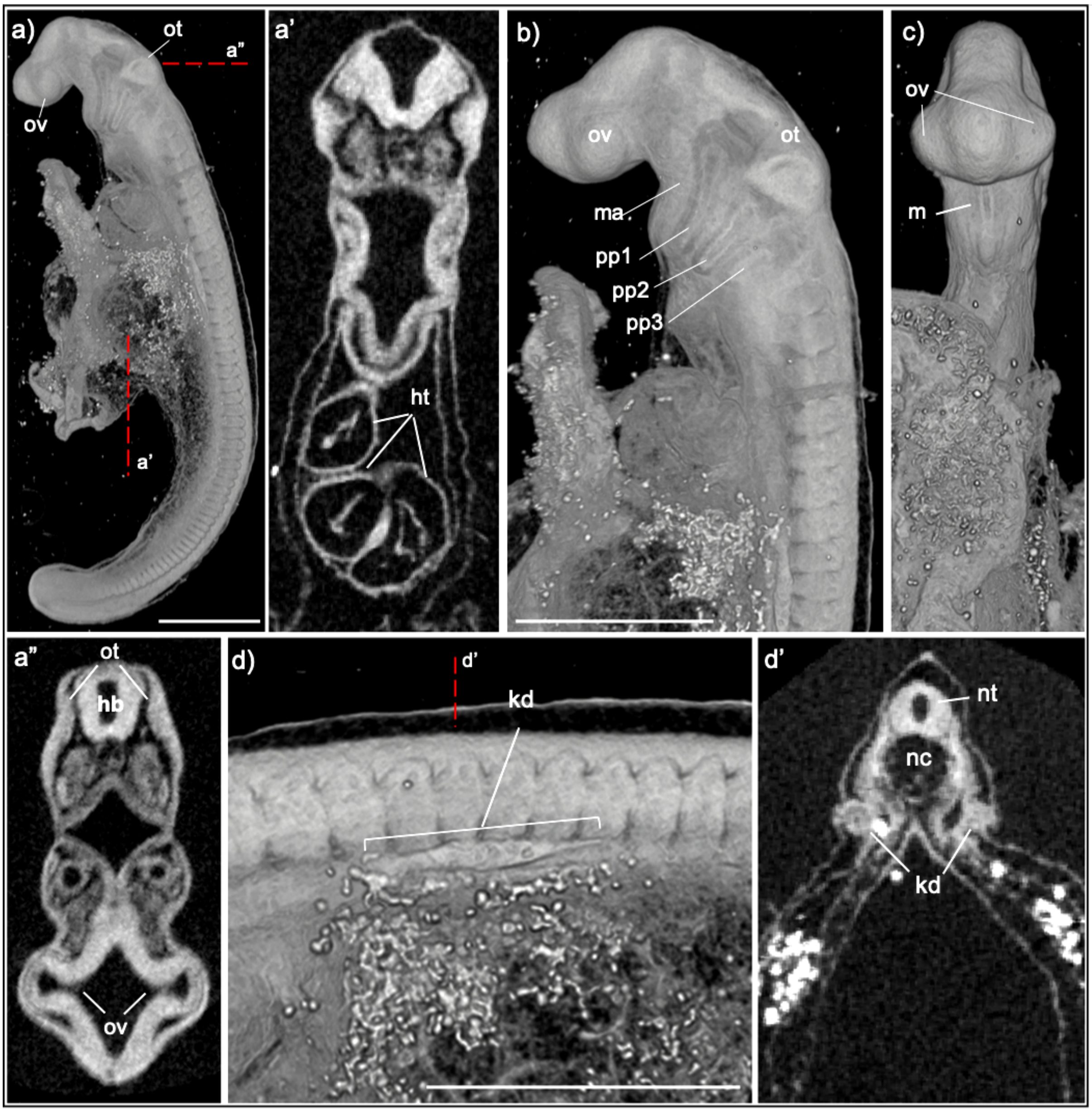


**Supplementary Figure 3. MicroCT of Stage 19 Epaulette Shark Embryo.** a. Left lateral view of a stage 19 epaulette shark embryo in whole mount. By this stage the otic placode (ot) is visible as a pit located at the back of the head, and when viewed in rendered cross section forms a distinct thickening on either side of the hind brain (hb) (a"). The developing optic vesicle (ov) has gone from a small bulge to a more pronounced out pocket of the fore brain, that has begun closing over to form a distinct vesicle (a"). a'. A rendered coronal section through the developing heart (ht) shows that the heart tube has begun the process of looping to form the distinct chambers of the adult heart. b-c. Left lateral (b) and ventral (c) views of the head show the prominent bulge of the optic vesicle and the otic placode pit. There is a deep indentation where the mouth (m) will form, which is framed by the mandibular arches (ma) on either side. Pharyngeal pouches 1, 2 and 3 are visible, but none are open at this stage. d. Left lateral view of the trunk shows the kidney duct (kd) along-side somite 6 to 11. d'. Rendered cross section through the site of the kidney duct. nt, the neural tube; nc, notochord. Scale bar 500μm.

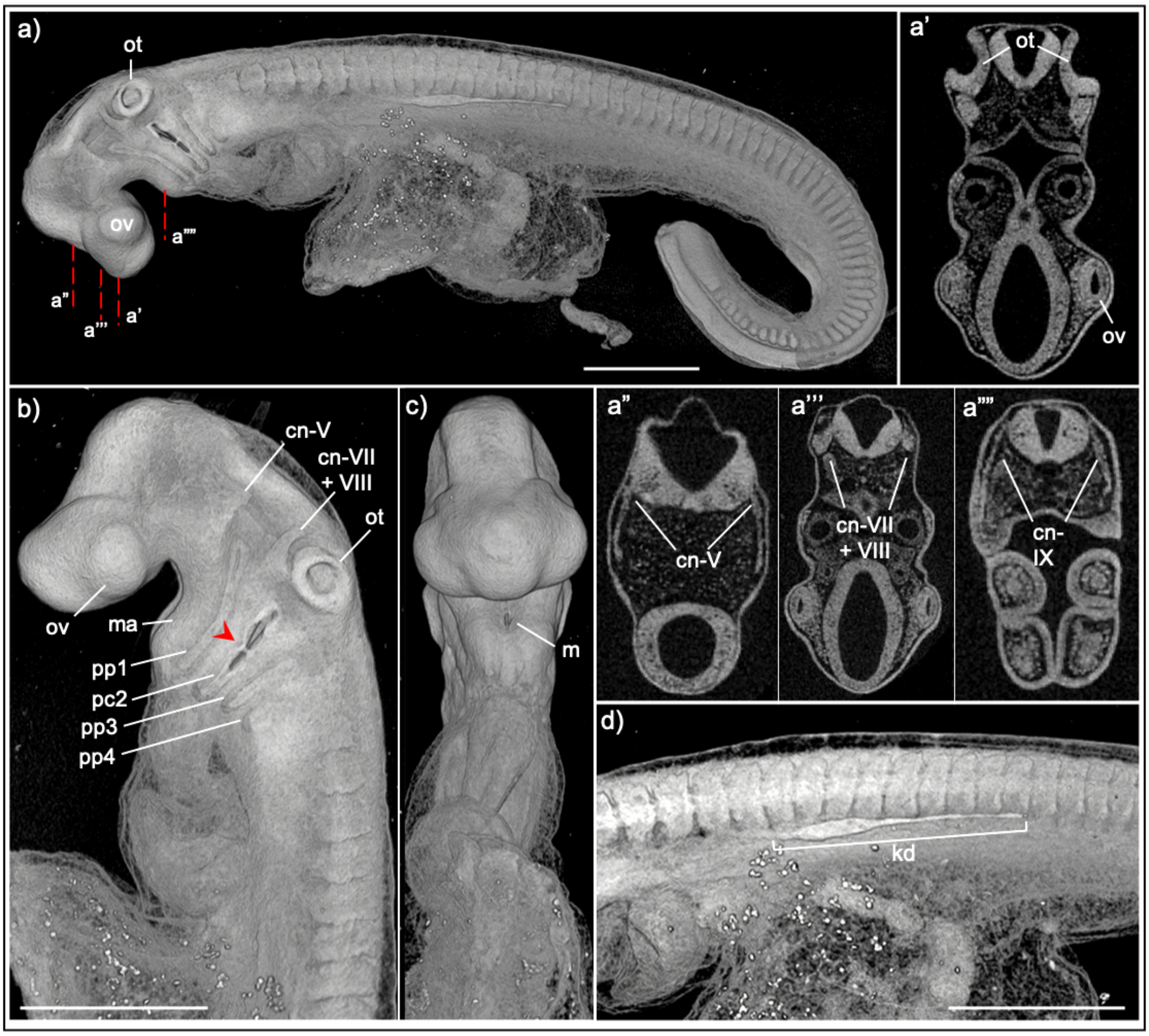


**Supplementary Figure 4. MicroCT of Stage 20 Epaulette Shark Embryo**. a. Left lateral view of a stage 20 epaulette shark embryo in whole mount, and rendered cross sections. **b-c.** Left lateral (b) and ventral (c) view of the head. The otic placode (ot) is a distinct pit clearly visible in whole mount (a,b) and cross section (a'). In rendered cross section and whole mount, several of the cranial nerves (cn-) are discernable, including cn-V, cn-VII + VIII and cn-IX (a'-a'''', b). Pharyngeal pouch 4 (pp4) is present by stage 20, and pharyngeal cleft 2 (pc2) has opened. Note that the formation of the cleft does not occur as a continuous opening, but appears to initiate in more than one location (red arrow). A small slit has started opening in the area where the mouth (m) will form (c). d. Left lateral view of the trunk shows that the kidney ducts (kd) extend further along the trunk by this stage and on the left side is visible along somites 7-16, whereas on the right hand side extends two somites further (i.e., from somites 7-18). Scale bar 500µm.

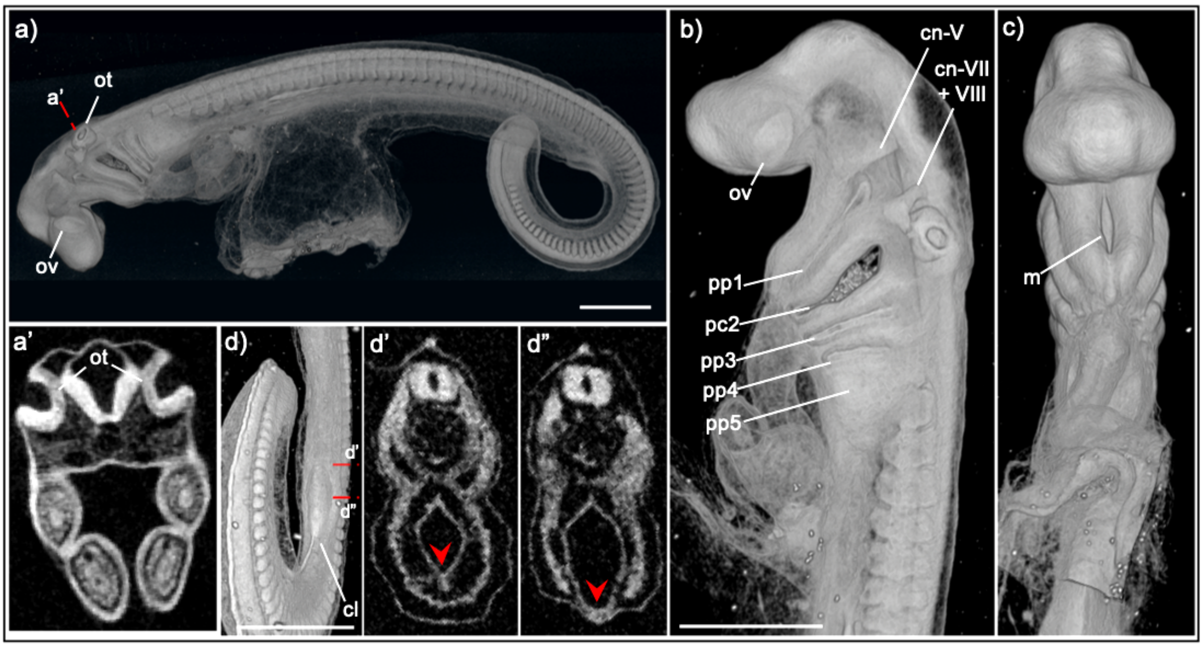


**Supplementary Figure 5. MicroCT of Stage 21 Epaulette Shark Embryo**. a-c. Left lateral view of a stage 21 Epaulette shark embryo in whole mount (a), left lateral (b) and ventral (c) view of the head. At stage 21, the otic pit (ot) has deepened (a'). b. Pharyngeal pouch (pp) 5 is visible as a slight swelling, and pharyngeal cleft 2 (pc2) has continued to open. In this specimen, cranial nerves (cn-) V and VII + VIII are prominent in lateral view. c. The slit where the mouth (m) will form is clearly visible in ventral view. d. Ventral view of the posterior trunk shows the position of the cloaca (cl) at the approximate axial level of somite 35-41. d'-d"The rendered cross section shows the changes in the body wall where the cloaca will form (red arrowhead). Scale bar 500μm.

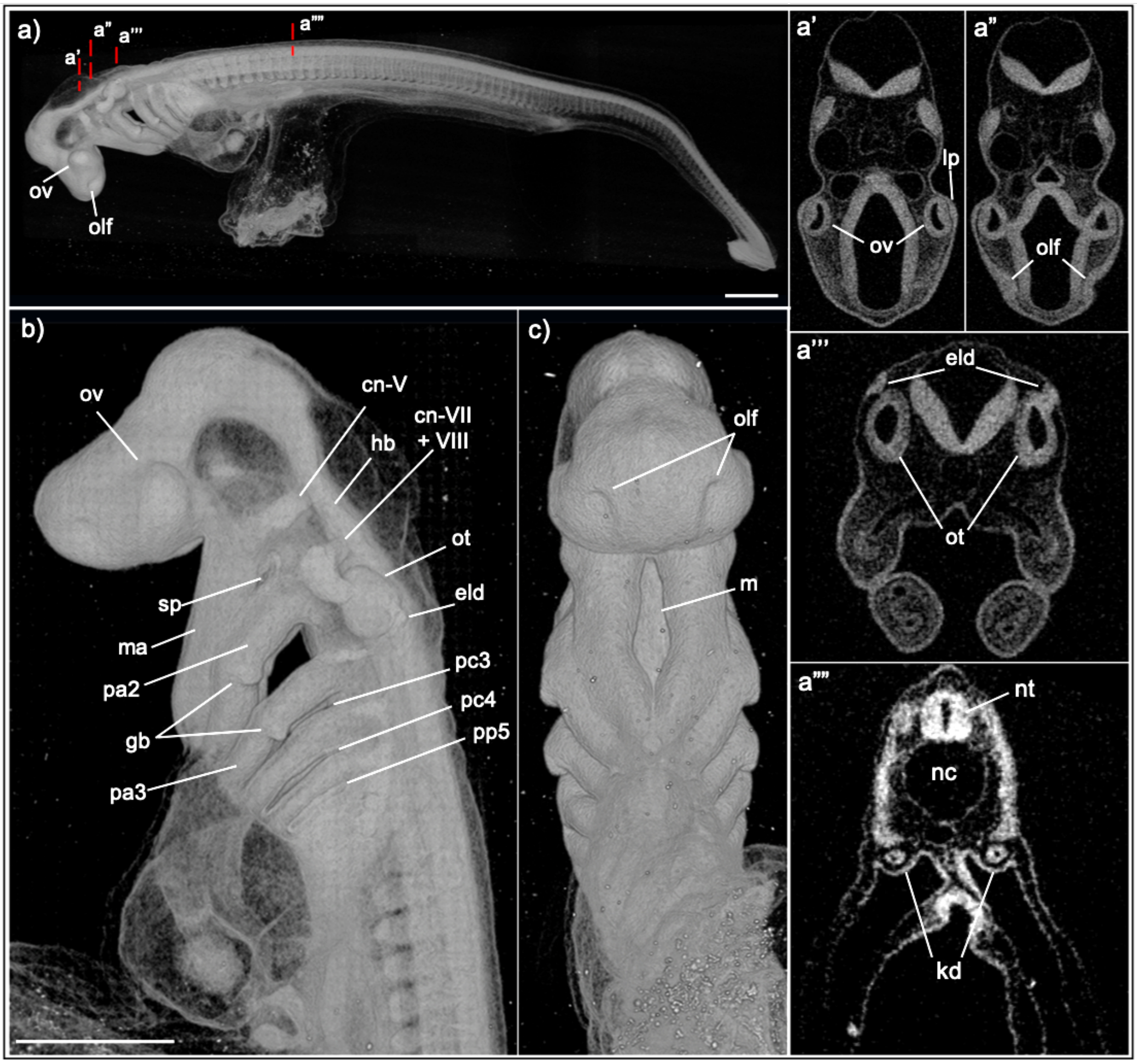


**Supplementary Figure 6.** MicroCT of Stage 22 Epaulette Shark Embryo. a. Left lateral view of a stage 22 Epaulette shark embryo in whole mount, and rendered cross sections. b-c. Left lateral (b) and ventral (c) view of the head. The lens placode (lp) forms as a thickening of the overlying ectoderm (a'). The olfactory pits (olf) appear as a thickening and slight indentation at the anterior tip of the head (a''). a''' The otic pit has now formed the enclosed otic capsule and the endolymphatic duct (eld) is visible extending towards the skin on the dorsal edge of the otic capsule. a'''' The kidney duct (kd) extends along the dorsal wall of the coelomic cavity. b The position of cranial nerves V, VII and VIII (cn-V, VII, VIII) remain visible in lateral view. Pharyngeal cleft 3 and 4 (pc3-4) are open, and pharyngeal pouch 5 (pp5) is visible. Pharyngeal arch 2 and 3 (pa2-3) bear a single gill bud (gb). nt, the neural tube; nc, notochord. Sp, spiracle. Scale bar 500µm.

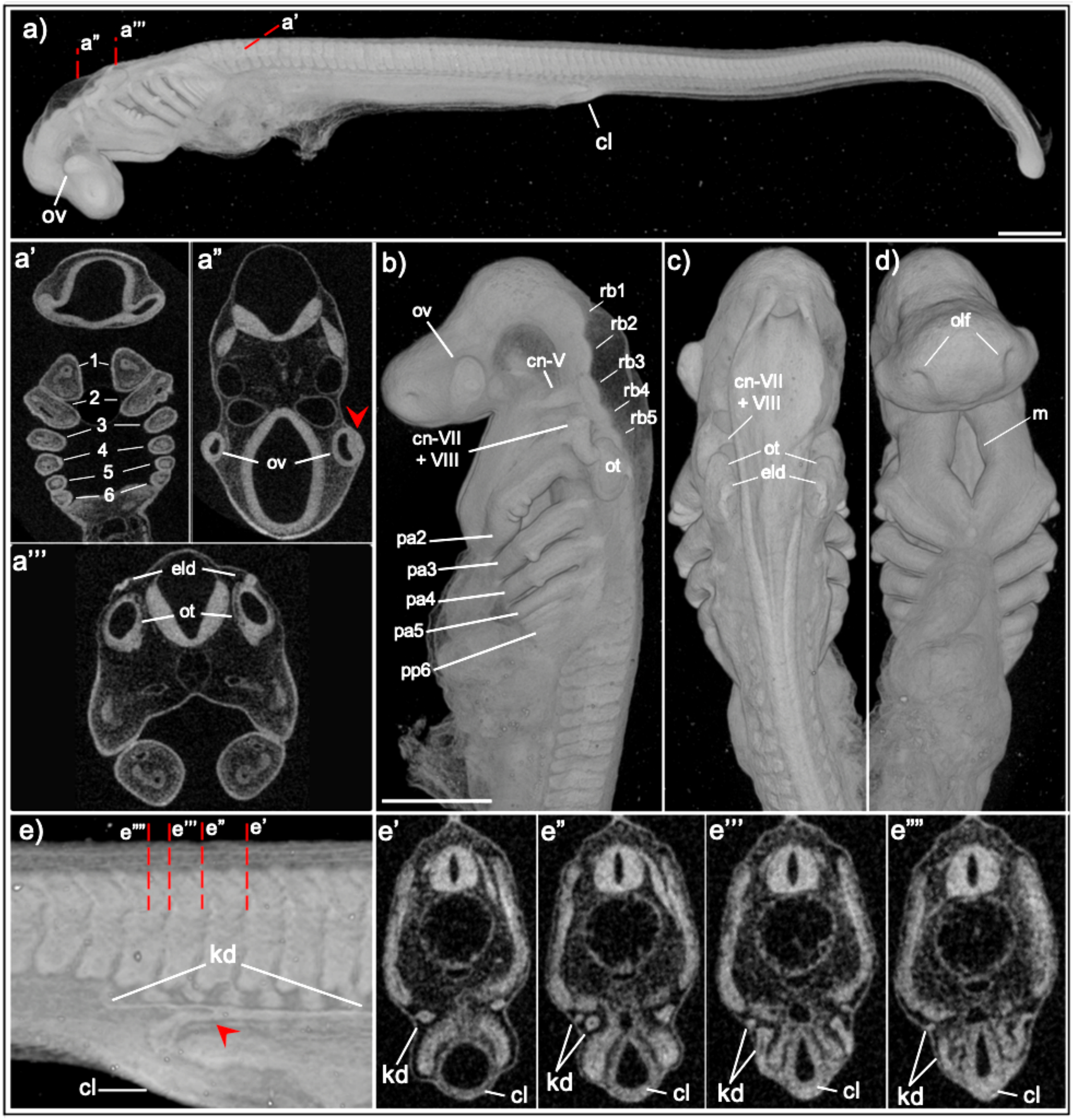


**Supplementary Figure 7. MicroCT of Stage 24 Epaulette Shark Embryo.** a. Left lateral view of a stage 24 epaulette shark embryo in whole mount and rendered sections. **b-d.** Left lateral (b), dorsal (c) and ventral (d) views of the head. By stage 24 there are 6 paired pharyngeal arches (pa) which can be seen in rendered section (a'). Three gill buds (gb) are present on pa2, and a single gill bud present on pa3 and pa4. Rendered cross section through the head show that the optic vesicle (ov) and lens placode (red arrow) are similar to the previous stage (a"), however, the concave shape of the optic vesicle is more pronounced (a''). The endolymphatic duct (eld) opens posteriorly from the otic capsule (ot) (a''', b). In this specimen, the rhombomeres (rb1-5) of the hindbrain are prominent in lateral view (b), as are cranial nerves (cn-) V and cn-VII + VIII. The mouth (m) slit is more open and has begun to widen, creating a diamond shape (d). e. Right lateral view of the body wall at the axial level of the cloaca (cl), anterior to right, shows the kidney duct (kd) splitting to form two distinct branches (red arrow), one extending ventrally towards the cl, and another branch extending posteriorly beyond the cl. e'-e"" Rendered cross sections through the posterior kd show a single duct (e’), splitting into two (e”), one extending ventrally towards the cl, whilst the other maintains its position ventral to the somites (e”’-e””). Scale bar 500µm.

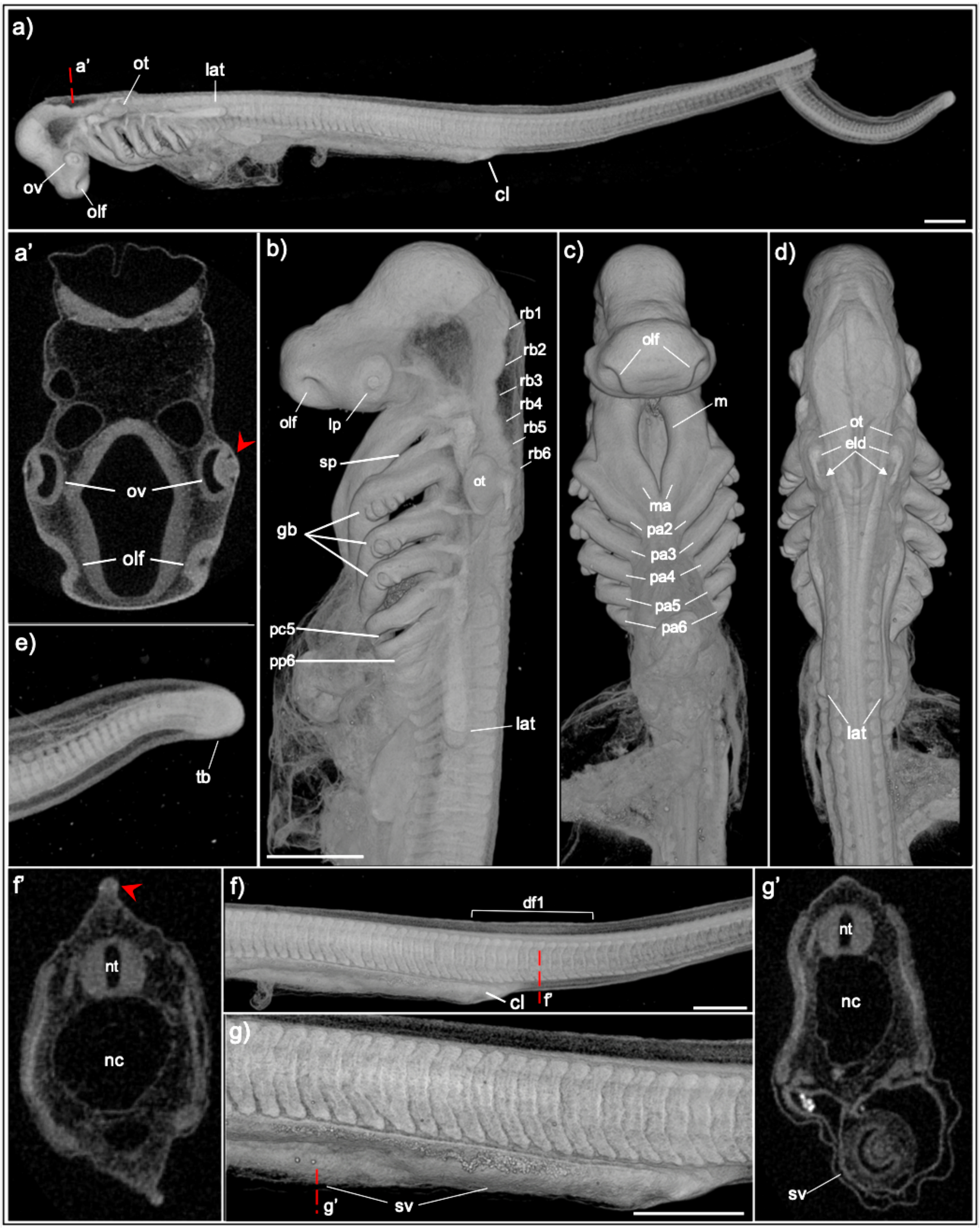


**Supplementary Figure 8. MicroCT of Stage 25 Epaulette Shark Embryo**. **a.** Left lateral view of a entire stage 25 epaulette shark embryo in whole mount, and rendered cross section (a'). **b-d.** Left lateral (b), ventral (c) and dorsal (d) views of the head. The position of the lens placode (lp) is now clearly visible in lateral view (b). The olfactory pit (olf) has become more pronounced (b,c). The spiracle (sp) and pharyngeal cleft 5 (pc5) are now open (b), pharyngeal pouch 6 (pp6) is visible as a slit, but not yet open (b, c). Gill buds (gb) are present on pa2 ($n$ = 5), pa3 ($n$ = 3) and pa4 ($n$ = 3) (b). At this stage the lateral line (lat) is visible branching from the post-otic ganglion, extending along the trunk to the axial level of somite 8 (b). There is a slight asymmetry to the lateral line progression along the trunk, with the right lateral line extending half a somite further than the left (d). **e**. The tail bud remains present at this stage (tb). **f-g.** Left lateral view of the trunk and rendered cross sections at select axial levels (indicated by red dashed line). Near the axial level of the cloaca (cl) there is a thickening of the median fin fold where the first dorsal fin (df1) will form (f'). In the posterior gut tube, the spiral valve (sv) has begun to form (g'). Scale bar 500μm.

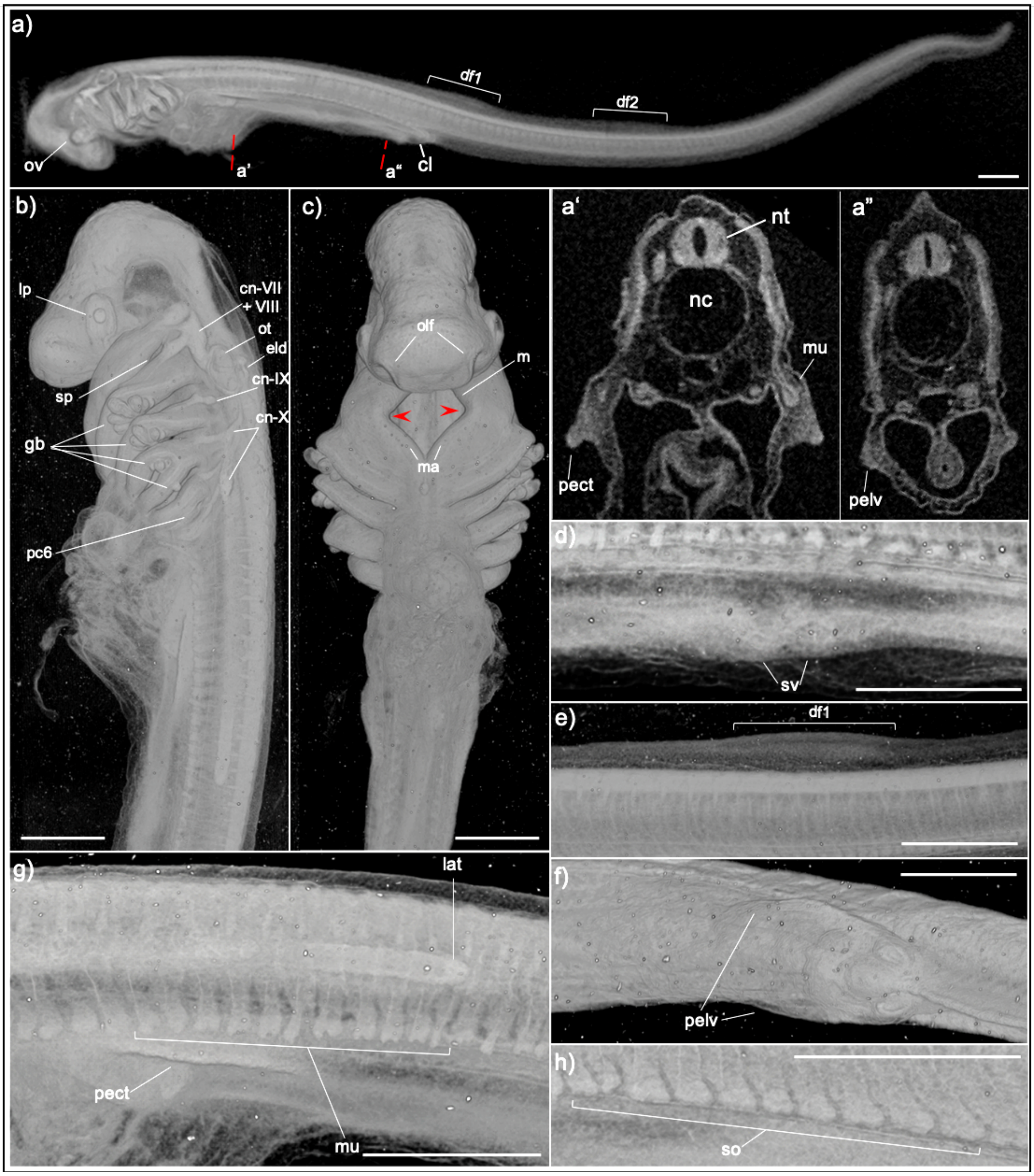


**Supplementary Figure 9. MicroCT of Stage 26 Epaulette Shark Embryo**. **a.** Left lateral view of a stage 26 epaulette shark embryo in whole mount, and rendered cross sections. **b-c.** Left lateral (b) and ventral (c) view of the head. **d-g.** Left lateral (d,e,g,h) and ventral (f) view of the trunk. The pectoral fins (pect) are visible by this stage as pronounced ridges along the trunk (a',g) with muscle buds (mu) beginning to extend from the ventral edge of the somite, toward the fin (a',g). Somites (so) posterior to the pectoral fins do not have the muscle bud extensions at this stage (h). The pelvic fins (pelv) are also visible as small ridges forming either side of the cloaca (cl) (a",f). Dorsal fin 1 (df1) and 2 (df2) are visible as slightly larger thickenings of the median fin fold (a,e). The mouth (m) begins to take on a pentagonal shape (c, red arrowhead). The endolymphatic duct (eld) has increased in length and extends caudal to the posterior margin of the otic capsule (ot). The lateral line (lat)

has extended further along the trunk, to the level of somite 20 (g). The spiracle (sp) and pharyngeal cleft 2-5 (pc2-5) are open, pharyngeal cleft 6 (pc6) is partially open (b-c). Gill buds (gb) are present on pa2 ($n$ = 7), pa3 ($n$ = 5), pa4 ($n$ = 3), pa5 ($n$ = 1). nt, neural tube; nc, notochord. Scale bar 500μm.

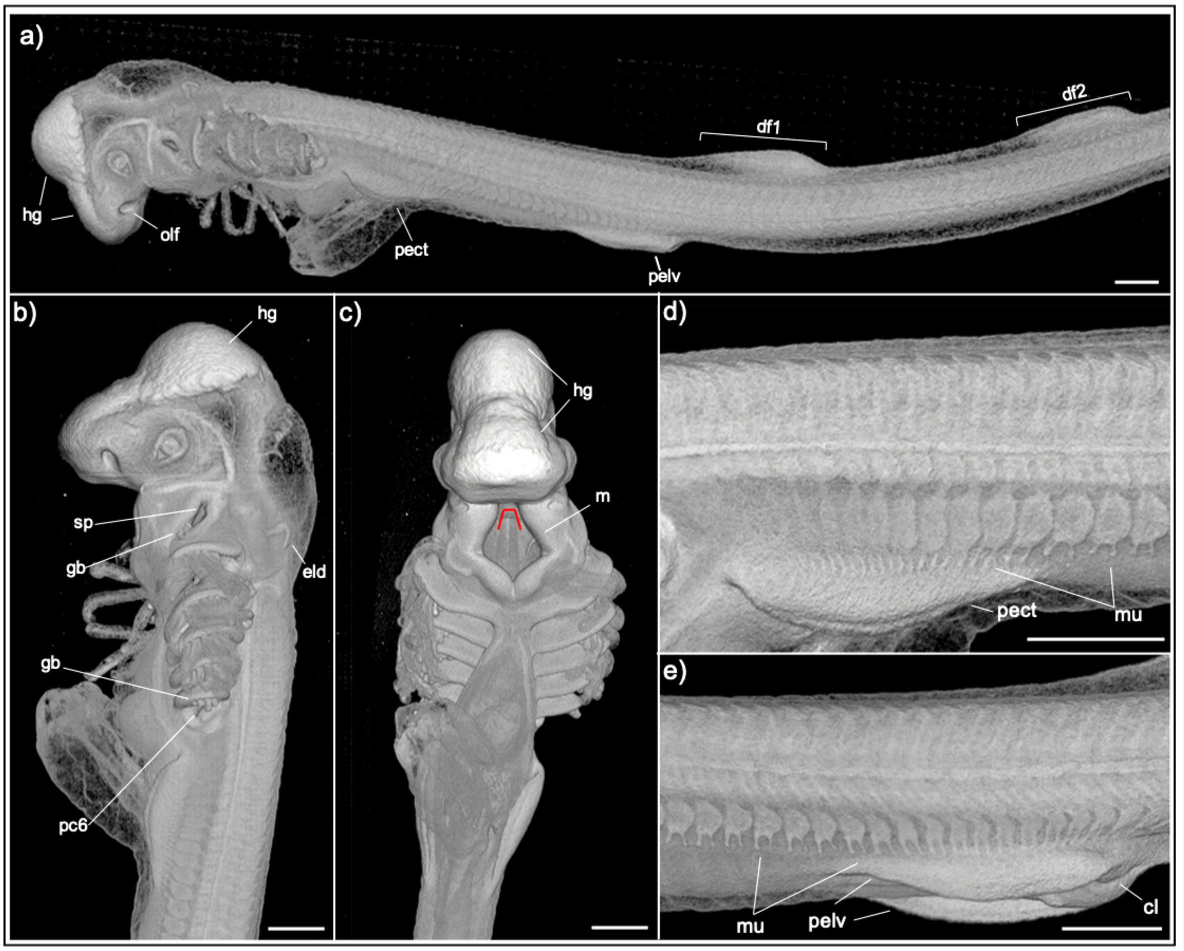


**Supplementary Figure 10. MicroCT of Early Stage 27 Epaulette Shark Embryo.**
**a.** Left lateral view of an early stage 27 epaulette shark embryo from the head to the second dorsal fin. **b-c.** Left lateral (b) and ventral (c) view of the head. **d-e.** Left lateral view of the trunk at the axial level of the pectoral fin (d) and pelvic fin (e). The hatching gland (hg) is visible as an external layer over the forebrain and midbrain region of the head (a-c). The first and second dorsal fins (df1, df2) are distinct outgrowths from the median fin fold, and the anal fin (af) is visible as a thickening of the ventral median fin fold (a). Pharyngeal cleft 6 (pc6) is fully open and gill buds (gb) are present on the spiracle (sp) and pharyngeal arch 2-6 (pa2-6). The mouth (m) is pentagonal in shape (c, red line). There are two distinct muscle buds (mu) extending from the ventral margins of each somite (mu) including at interfin levels. At pectoral and pelvic levels, paired ventral muscle buds extend to the base of the fins.
Scale bar 500μm.

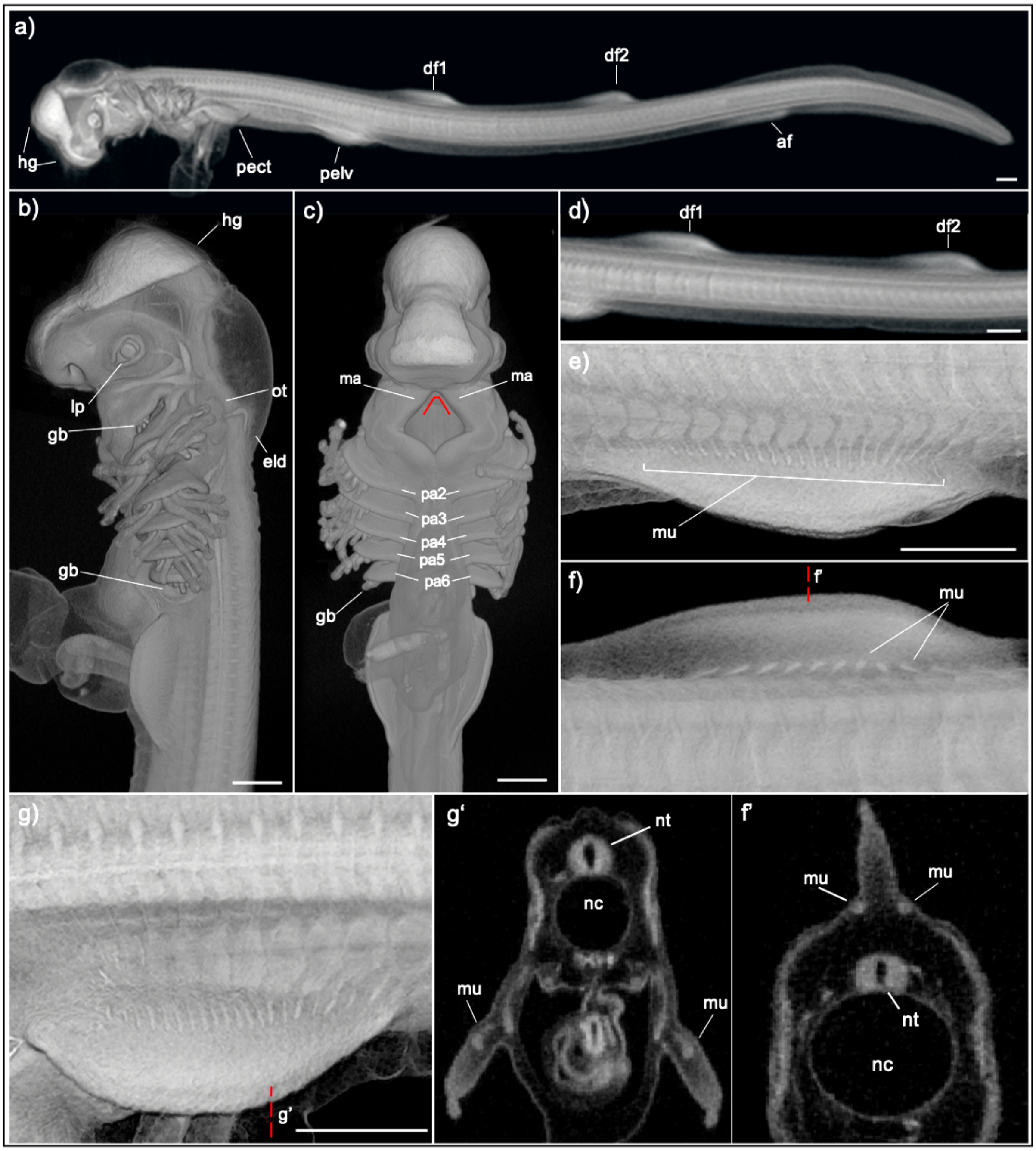


**Supplementary Figure 11**. **MicroCT of Later Stage 27 Epaulette Shark Embryo.** **a.** Left lateral view of a later stage 27 epaulette shark embryo in whole mount. **b-c.** Left lateral (b) and ventral (c) view of the head. **d-g.** Left lateral view of the trunk and rendered cross sections. The hatching gland (hg) remains visible over the forebrain and midbrain region of the head (a-c). The first and second dorsal fins (df1, df2) are distinct outgrowths from the median fin fold (a). Gill buds (gb) are present on the spiracle (sp) and pharyngeal arch 2-6 (pa2-6). The mouth remains slightly pentagonal in shape (red line), though the space between maxillary processes is reduced. The paired muscle buds (mu) extend ventrally into the pectoral (g) and pelvic fins, as well as along the body wall at interfin levels. A single projection extends from each somite neighbouring the dorsal fin towards the fin base (f, f') nt, neural tube; nc, notochord. Scale bar 500μm.

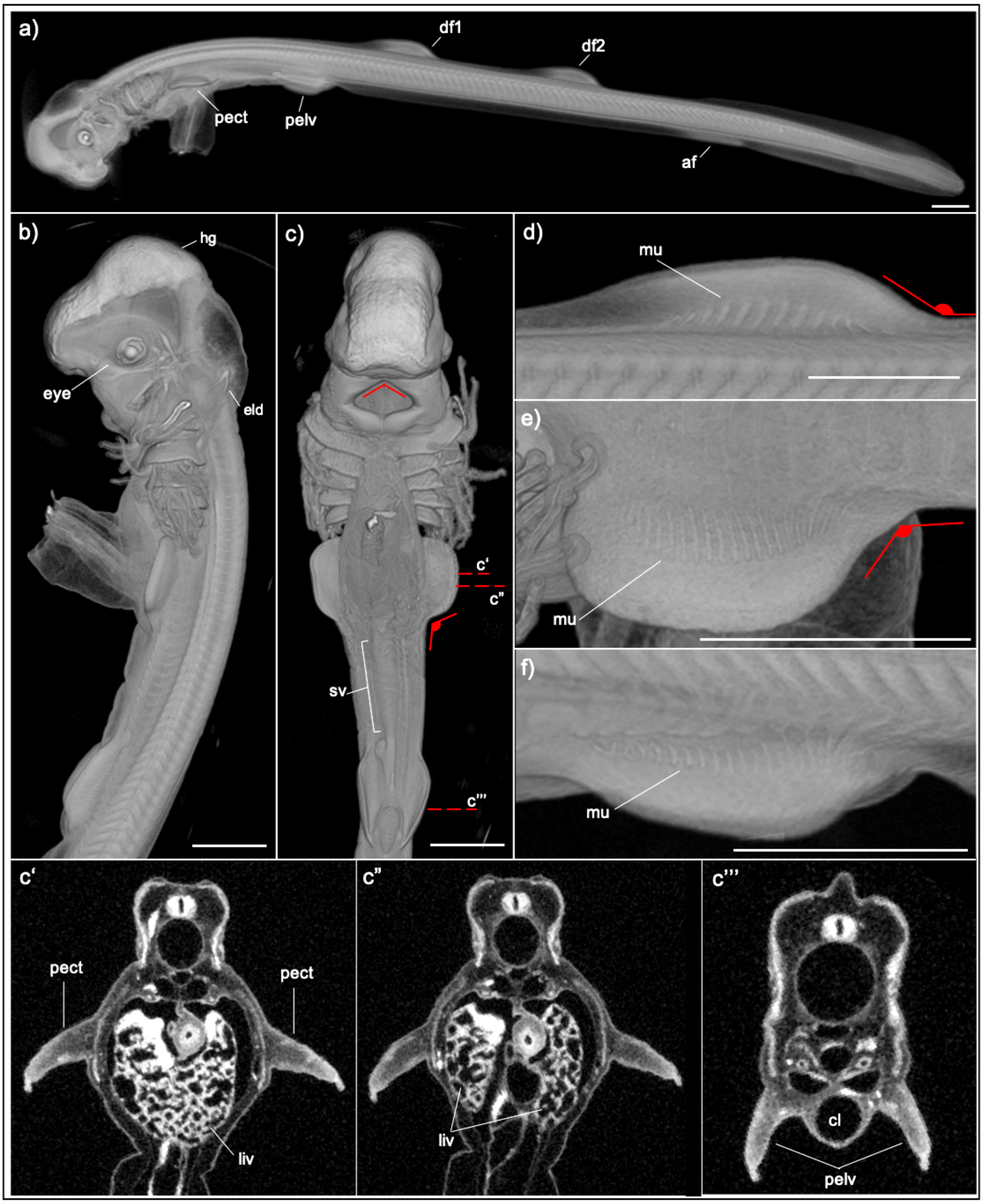


**Supplementary Figure 12. MicroCT of Stage 28 Epaulette Shark Embryo. a.** Left lateral view of a stage 28 epaulette shark embryo in whole mount. **b-c.** Left lateral (b) and ventral (c) view of the head and rendered cross sections. **d-f.** Left lateral view of the trunk at the axial level of dorsal fin 1 (d), the pectoral fin (e), and pelvic fin (f). The hatching gland (hg) is visible over the region of the forebrain and midbrain (b,c). The mouth is diamond shaped (red line) (c). The posterior margin of dorsal fin 1 and 2 (df1, df2) form an obtuse angle with the trunk (d). The pectoral fin forms an obtuse angle with the body (c,d), and muscle buds can be seen extending further distally into both the pectoral (e) and pelvic fins (f). Rendered cross sections through the

Level of the pectoral fin show the continuous liver parenchyma (liv) anteriorly (c’) and two separate lobes more caudally (c”). c”’.Rendered cross section through the pelvic fins also shows the unperforated cloaca. sv, spiral valve. Scale bar 1mm.

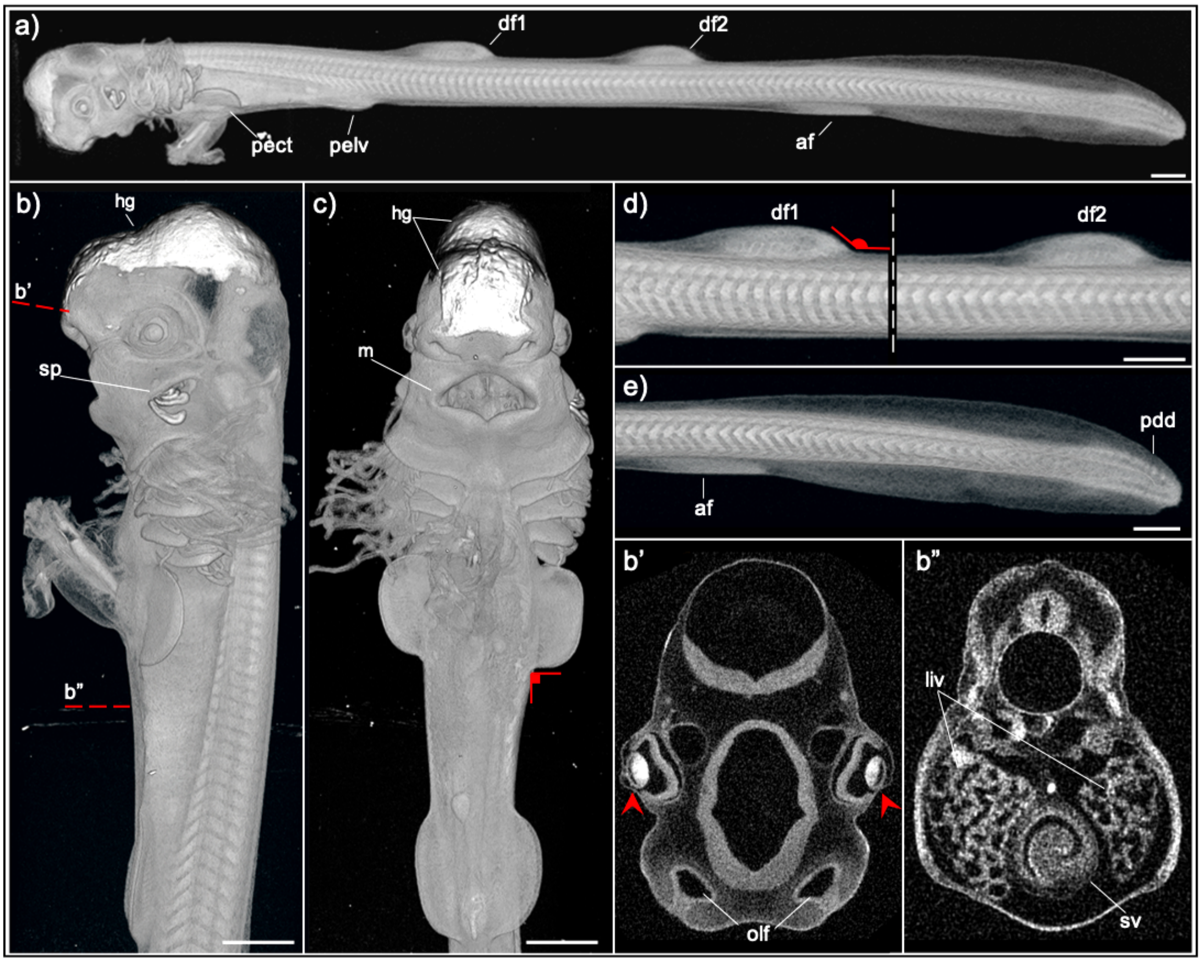


**Supplementary Figure 13**. **MicroCT of Stage 29 Epaulette Shark Embryo.** MicroCT of Stage 29 Epaulette Shark Embryo. **a.** Left lateral view of a stage 29 epaulette shark embryo in whole mount. **b,c.** Left lateral (b) and ventral (c) view of the head to the pelvic fins, and rendered sections. **d,e**. Left lateral view of the trunk. The dorsal fins 1 and 2 (df1, df2) form an obtuse angles with the body (a,d), and the anal fin (af) is visible as a thickening of the ventral median fin fold that has begun to extend distally from the surrounding tissue (a,e). The primordia of dermal denticles (pdd) are visible at the caudal end of the tail (e). The pectoral (pect) and pelvic (pelv) fins continue to grow larger (a,c), and the pect fin now forms a right angle with the body in ventral view (c). The olfactory pits extend further into the surrounding tissue to now form a deep olfactory chamber, visible in rendered cross section (b'). The eyes continue to develop and the lens is no longer in contact with the underlying optic vesicle (b'). b" Rendered cross section midway down the trunk shows two lobes of the developing liver (liv), and characteristic looping of the spiral valve (sv). hg, hatching gland; m, mouth. Scale bar 1mm.

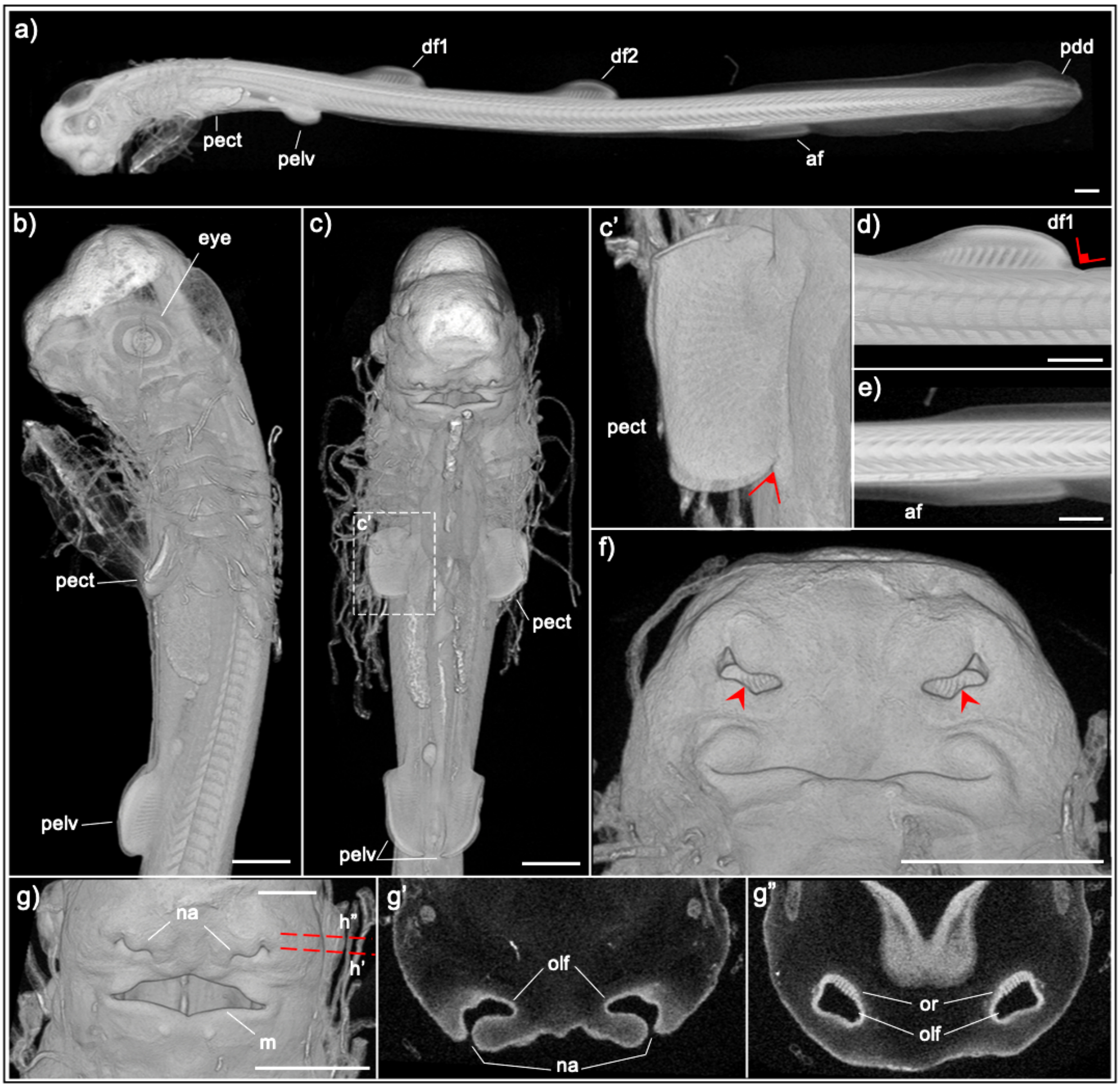


**Supplementary Figure 14**. **MicroCT of Stage 30 Epaulette Shark**. **a.** Left lateral view of a stage 30 epaulette shark embryo in whole mount. **b,c.** Left lateral (b) and ventral (c) view of the head and trunk. **d,e**. Left lateral view of the trunk. **f,g.** Ventral view of the head and rendered sections. The pectoral (pect) fins form an acute angle with the body (c'), whereas dorsal fins 1 and 2 (df1, df2) form a right angle with the body (a,d). The anal fin (af) begins to extend distally from the surrounding fin fold (a,e). The external opening of the olfactory chamber (olf) begins to narrow and constrict in the middle, defining the positions of the external nares medially and laterally (f). In rendered cross section, the olfactory rosettes (or) are visible as distinct parallel grooves forming at the back of the olfactory chamber (g, g"). Scale bar 1mm.

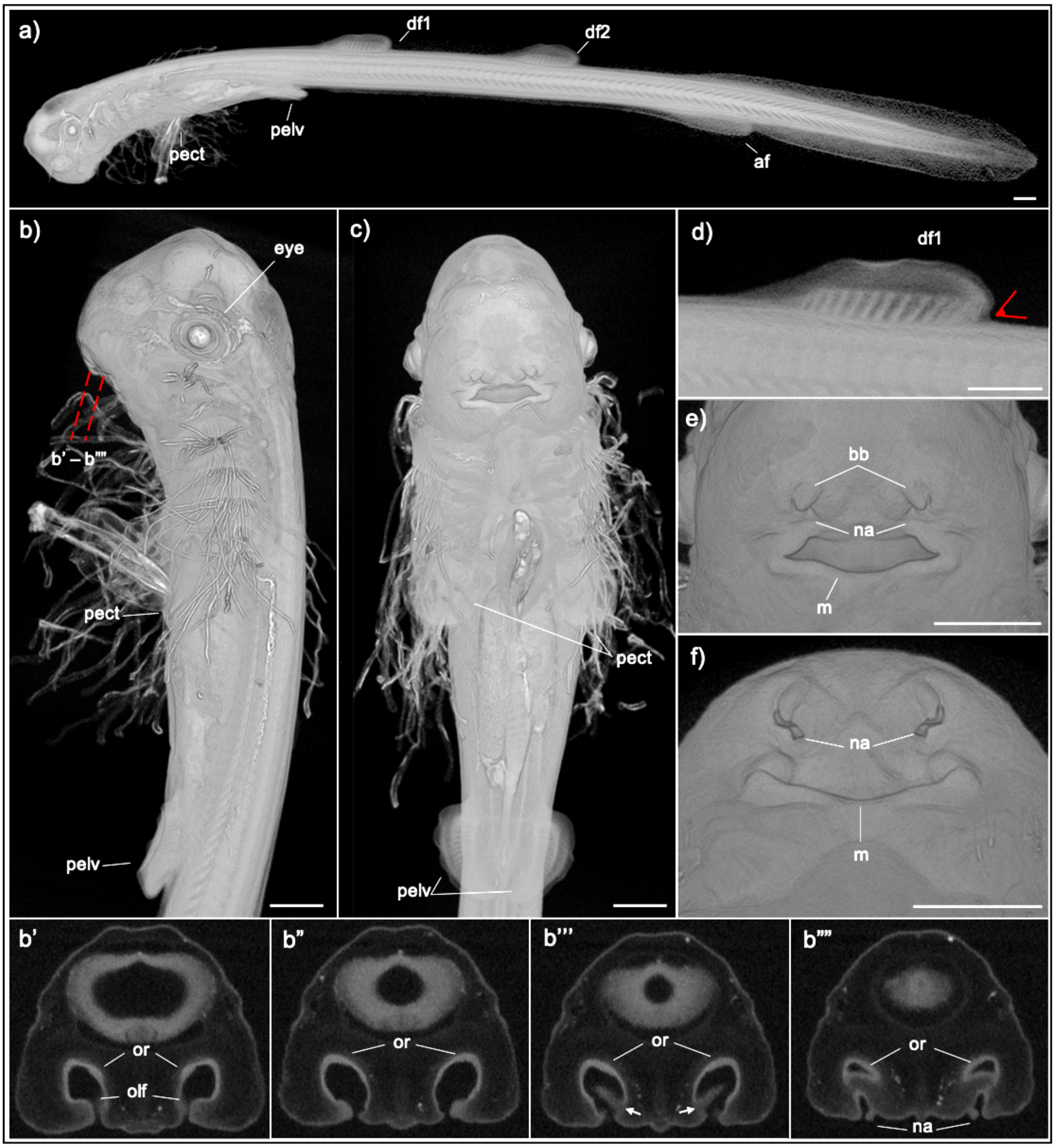


**Supplementary Figure 15**. **MicroCT of Stage 31 Epaulette Shark** a. Left lateral view of a stage 31 epaulette shark in whole mount. **b,c.** Left lateral (b) and ventral (c) view of the head and trunk and rendered sections. **d.** Left lateral view of the trunk. **e,f.** Ventral view of the head. The dorsal fins 1 and 2 (df1, df2) form a slightly acute angle with the body (a,d). The nares (na) continue to narrow, and the surrounding tissue begins to form distinct protrusions, which form the barbels (bb) (e,f). The olfactory chambers (olf) have distinct folds (arrow) beginning to form, directing water across the olfactory rosettes (or) located at the back of the chambers (b'-b''''). af, anal fin; m, mouth. Scale bar 1mm.
